# The Oligomerization Landscape of Histones

**DOI:** 10.1101/422360

**Authors:** Haiqing Zhao, David Winogradoff, Yamini Dalal, Garegin A. Papoian

## Abstract

In eukaryotes, DNA is packaged within nucleosomes. The DNA of each nucleosome is typically centered around an octameric histone protein core: one central tetramer plus two separate dimers. Studying the assembly mechanisms of histones is essential for understanding the dynamics of entire nucleosomes and higher-order DNA packaging. Here we investigate canonical histone assembly and that of the centromere-specific histone variant CENP-A using molecular dynamics simulations. We quantitatively characterize their thermodynamical and dynamical features, showing that two H3/H4 dimers form a structurally floppy, weakly-bound complex, the latter exhibiting large instability around the central interface manifested via a swiveling motion of two halves. This finding is consistent with the recently observed DNA handedness flipping of the tetrasome. In contrast, the variant CENP-A encodes distinctive stability to its tetramer with a rigid but twisted interface compared to the crystal structure, implying diverse structural possibilities of the histone variant. Interestingly, the observed tetramer dynamics alter significantly and appear to reach a new balance when H2A/H2B dimers are present. Furthermore, we found that the preferred structure for the (CENP-A/H4)_2_ tetramer is incongruent with the octameric structure, explaining many of the unusual dynamical behaviors of the CENP-A nucleosome. In all, these data reveal key mechanistic insights and structural details for the assembly of canonical and variant histone tetramers and octamers, providing theoretical quantifications and physical interpretations for longstanding and recent experimental observations. Based on these findings, we propose different chaperone-assisted binding and nucleosome assembly mechanisms for the canonical and CENP-A histone oligomers.

## INTRODUCTION

Eukaryotes wrap their DNA around histone proteins constituting the fundamental unit of chromatin, the nucleosome. Inside each nucleosome, histones typically exist as an octamer, composed of a central tetramer (H3/H4)_2_ plus one H2A/H2B dimer on either side (1). Nucleosomes dynamically dissociate and re-associate in chromatin structure for fundamental biological processes such as DNA transcription, replication, and repair. By initiating nucleosome assembly through forming a tetrasome with DNA, the histone tetramer serves as the structural basis for nucleosomal or chromatin dynamics (2, 3). Thus, it is crucial to elucidate the dynamics of histone tetramers, which are key intermediates along nucleosome assembly and disassembly pathways. Recent single-molecule experiments studied the spontaneous flipping behavior of DNA handedness of the tetrasome, finding that iodoacetamide-treated residue mutations on the tetramer can cause the enhanced flexibility and faster superhelical flipping kinetics of the wrapped DNA (4–6). Therefore, a deep molecular understanding of histone tetramer dynamics is not only essential to understanding subnucleosomal or nucleosomal assembly, but may also suggest innovative pathways for higher-order DNA packaging.

The centromere-specific histone H3 variant, Centromere Protein A (CENP-A), has been extensively studied in recent decades: (1) for its significant functional role as the epigenetic mark of centromere ensuring proper chromosome separation during cell division (7–12); (2) for its unique structural dynamics (13–15), particularly dissecting the dominant structure of CENP-A nucleosomes (16–21) and their special association with kinetochore partners (22–26). Unlike canonical H3 nucleosomes, CENP-A-containing nucleosomes follow a different assembly pathway *via* the unique chaperone HJURP (27–31). Also, in cancer cells CENP-A is over-expressed and the redundant CENP-A can localize into ectopic (*i.e.* non-centromeric) regions *via* alternative pathways (32, 33). Thus, one outstanding question is whether CENP-A, in normal cells, can be efficiently regulated to avoid ectopic delivery. Another related question is whether replacing canonical H3 with CENP-A alters its physical properties and overall dynamics.

Conflicting studies have suggested that: (1) *in vitro* chromatography and deuterium exchange experiments indicate that the soluble CENP-A/H4 forms a more compact and rigid tetramer than the conventional H3 complex (34); partially truncated CENP-A tetramers adopt compact conformations in crystals and in solution (16); (2) CENP-A- and H3-containing nucleosomes have nearly identical crystal structures (35, 36), and (3) recent computational and experimental studies reveal that CENP-A dimers (37) and nucleosomes (38, 39) are more flexible than their canonical H3 counterparts. On the other hand, canonical histone tetramers present consistent crystal structures in different molecular contexts, including as a tetramer in a nucleosome (1,40), in an octamer (41–43), and in complexes with chaperones such as FACT (44), Spt2 (45), TONSL and MCM2 (46, 47). Early size-exclusion chromatography experiments demonstrate that there is a dynamic equilibrium between two H3/H4 dimers and an assembled tetramer (48, 49), and this equilibrium is responsive to changes in ionic strength (50). Through Electron Paramagnetic Resonance (EPR) spectroscopy, a previous study shows that canonical histone tetramer exhibits greater structurally heterogeneity on its own than when sequestered in the octamer (51). However, dynamical structural details that would reveal the mechanisms governing observed properties are not readily amenable to existing experimental techniques. Here we apply computational modeling to study both H3- and CENP-A-oligomers to provide a comprehensive theoretical quantification that can explain and unify these experimental observations that might seem incompatible.

Among computational approaches, molecular dynamics (MD) simulations are able to capture mechanistic details at the molecular level, complementing experimental approaches. Previously, we used atomistic MD to reveal that the CENP-A nucleosome exhibits greater flexibility than the canonical nucleosome around their native states (38), and its dynamics can be modulated by internal modifications (52). Combining coarse-grained, atomistic simulations and *in vivo* mutation experiments, we reported that the CENP-A dimer is structurally variable, and chaperone HJURP prevents the promiscuous mis-assembly of the CENP-A dimer, protecting it from binding with other proteins (37).

Building upon these findings, we performed coarse-grained MD simulations using the Associative-memory, Water-mediated, Structure and Energy Model (AWSEM) model (53, 54) to investigate the assembly mechanisms of histone oligomers and asked whether histones CENP-A and H3 differ at the tetramer/octamer level. We computed the association free energy of two dimers forming a tetramer, finding that CENP-A dimers form a more compact and stable tetramer with more favorable free energy, while the landscape of H3 dimers is more rugged, indicating its structural lability. In particular, simulations starting from pre-assembled tetramers reveal swiveling motion around the H3 tetrameric interface. Furthermore, histone octamer simulations suggest that the addition of H2A/H2B dimers gently restrains the internal rotation of the H3 tetramer. In contrast, H2A/H2B addition causes the CENP-A tetramer to adopt multiple conformational states, demonstrating a significant incongruence between the preferred structures of the CENP-A tetramer versus the octamer. Finally, we put forward a speculative model for canonical and variant histone assembly and propose that the CENP-A tetramer may serve as a critical sequestration channel, preventing the assembly of excess CENP-A into chromatin, thereby regulating CENP-A homeostasis *in vivo*.

## MATERIALS AND METHODS

### Initial structures

In spite of diverse structural environments, the canonical histone tetramer adopts a consistent configuration in the crystal structures of histone octamer, nucleosome and protein complex with chaperone protein(s) (detailed comparisons are provided in Supporting Figure S4). In this work we took the tetramer structure from a nucleosome crystal structure containing H3 (PDB: 1KX5 (40)). Initial configurations for CENP-A tetramer were obtained from the CENP-A-containing nucleosome (PDB: 3AN2 (35)) and the *αN*-helices-truncated CENP-A tetramer crystal structure (PDB: 3NQJ (16)). Current study does not include histone tails and DNA. Their effects are discussed in the Discussion section. More structural information of the excluded nucleosomal DNA and histone tails (Figure S5), and information about simulated protein length and their sequences (Figure S6) are covered in the SI.

### Simulation methods and trajectory analyses

In this work, we used the Associative-memory, Water-mediated, Structure and Energy Model to carry out computational simulations for both canonical and variant CENP-A histone oligomers. AWSEM is a coarse-grained protein model with three beads (C_*α*_, C_*β*_ and O) representing one amino acid. The total potential function includes the *V_backbone_* term for protein backbone formation, residue-residue and residue-water interaction terms *V_contact_* and *V_burial_*, hydrogen bonding term *V_HB_*, and the associate memory term *V_AM_* (Eq. 1). Details of each potential term is described in the supplemental material of ref. 54 and the SI of this work. This model is based on not only the physical interactions like *V_backbone_* and *V_HB_* but also the bioinformatics-inspired fragment memory term *V_AM_* (Figure S1). Here we use the respective histone monomer structures to build the biasing structural fragment memory database wherein each fragment is 3- to 12-residue long. It is important to note that no intermolecular information between either monomers or dimers were provided to the force field. So, from this perspective, AWSEM is used as a predictive protein model. In the constant temperature simulations of two dimers, a weak distance constraint in the harmonic potential form (spring constant *k* = 0.02 kcal/mol/Å^2^) is applied between center of masses of each dimer. Constraints of the same magnitude are applied between the two H2A/H2B dimers, and between each H2A/H2B dimer and the tetramer. These constraints ensure the two objects are within a reasonable distance of each other for possible interactions.

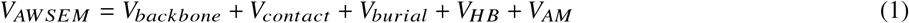

All simulations were performed in the large-scale atomic/molecular massively parallel simulator 2016 (LAMMPS 2016), using the Nosé-Hoover thermostat. We applied umbrella-sampling together with replica-exchange (55) to enhance the phase space sampling for further free energy calculations. For instance, in the case of H3 dimers’ association, two H3 dimers were put in the simulation with the distance between their centers-of-mass controlled by an umbrella constraint. A typical harmonic potential is used for this purpose as shown in Eq.(2), where *k_R_* is the biasing strength and *R_o_* is the controlled center distance for each window. Here *k_R_* = 5 kcal/mol/Å^2^, and *R_o_* ranges from 20 Å to 50 Å, averagely spaced by 1 Å. Simulations for each umbrella window, in total 30, were performed using ten replicas with temperatures linearly ranging from 280 K to 370 K. After the simulation reached convergence (see below and 4), data from different windows were then collected and the weighted histogram analysis method (WHAM) (56) was applied to calculate the PMFs and construct free-energy landscapes onto different coordinates. A relevant Jacobian factor correction term, *k_B_T* ln[4π*R*^2^], was subtracted from the free energy calculation since a sampling space based on the distance *R_COM_* makes nonphysical contributions to the configurational partition function (57). The time step is set at 5 fs in all simulations. Each replica was run for 2 million steps. Exchanges between replicas were attempted every 400 steps. The first 0.5 million steps were not used in the analysis for system’s equilibration.

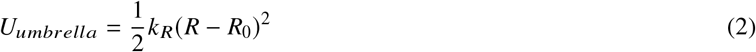

Separately, ten independent constant temperature simulations were carried out for tetramers (H3/H4)_2_ and (CENP-A/H4)_2_, with 30 million timesteps each and 300 million steps in total (1500 ns in the coarse-grained timescale). Weak biases in the form of harmonic potential were applied as mentioned above. Simulations and analyses for H3 and CENP-A tetramers that exclude *αN* helices are performed using the same setup. Octamer simulations for (H3/H4)_2_ and (CENP-A/H4)_2_ with two (H2A/H2B)s were run for 10 million timesteps, totaling 100 million timesteps for each octamer system. Simulations were performed in a 200-Å-long cubic box with periodic boundary conditions. Trajectories were combined for later data analysis after removing the first 10 ns in every run to account for thermal equilibration. Note that the coarse-graining timescale cannot be directly converted into real time since it could be at least 10 times larger than that in the atomistic MD simulations (58). The convergences of all simulations were verified by the root-mean-squared inner-product (RMSIP) analysis, which are provided in Supplemental Section 4.

All the trajectory analyses in this work, including the calculations of root-mean-square deviations (RMSD), radius-of-gyration (*R_g_*), distances (*R*), dihedral angles *θ, Q* values, and contact analysis, were based on the C*α* coordinates. More analyzing details including error analyses are included in Supplemental Sections 2 and 3.

## RESULTS

### Binding free energy of two dimers forming a tetramer

Motivated by the previous observation of CENP-A dimer flexibility (37) compared with its canonical counterpart, we first investigated the formation of tetramers from two canonical H3 and CENP-A dimers. *Via* a mixed enhanced sampling methodology that couples replica-exchange with umbrella-sampling, we mapped their corresponding binding free energy landscapes. The calculated free energy profiles (FEP) were projected into two reaction coordinates: the distance between centers-of-mass of the two dimers *R_COM_* and another order parameter *Q_interface_* which quantifies the nativeness of the binding interface between *Q_interface_* is the fraction of native interface contacts, defined as, 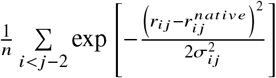, where *n* is the total number of contacts, *r_ij_* is the distance between the Cα atoms of residues *i* and *j*, and *σ_ij_* is given as *σ_ij_* = (1 + |*i − j*|^0.15^). *Q* ranges from 0 to 1, where no common contacts between a conformation and the native state corresponds to 0 and complete similarity of contacts corresponds to 1. Here the *Q_interface_* calculations were conducted with respect to the tetramer interface from the corresponding nucleosomes containing canonical H3 and variant CENP-A (PDB: 1KX5 (40) and 3AN2 (35)).

As seen in Figure 1, the binding free energy landscape for H3/H4 dimers is relatively rugged with multiple energy minima, at *Q_interface_* = 0.4, 0.1~0.2, and 0.0; *R_COM_* of 29 Å, 32~33 Å and 38 Å in the other dimension (Figure 1A). These minima are relatively flat compared to that of CENP-A, occupying a large portion of configuration space described in terms of *R_COM_* and *Q_interface_*, indicating larger structural heterogeneity of (H3/H4)_2_ with a broad ensemble of accessible conformations. This result is consistent with the experimental observation that histone H3 tetramer is unstable at moderate ionic strengths, determined by guanidinium-induced denaturation (48). On the other hand, the two CENP-A/H4 dimers present a deep, well-funneled association free energy landscape (Figure 1B), with the minimum at *R_COM_* = 29 Å, *Q_interface_* = 0.4, suggesting that there is a thermodynamically favorable binding state for the tetramer (CENP-A/H4)_2_.

**Figure 1:**
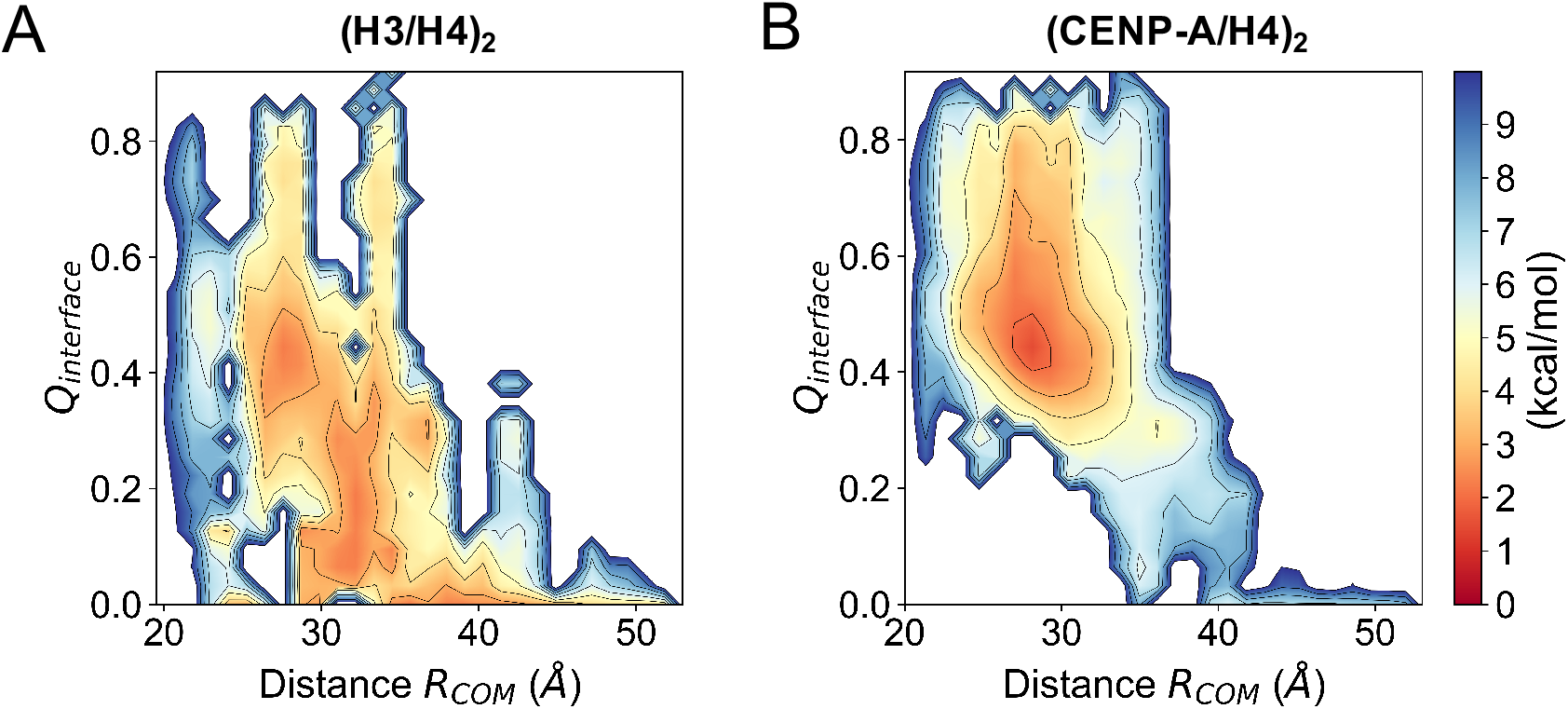
The binding free energy landscapes of two H3 dimers and that of two CENP-A dimers. Two-dimensional free energy profiles are mapped as a function of the distance between two interacting dimers *R_COM_* and of the quantification of the nativeness of their binding interface *Q_interface_*, for (H3/H4)_2_ (A) and (CENP-A/H4)_2_ (B).

To further quantitatively compare the binding energy differences between H3/H4 and CENP-A/H4, we projected two computed FEPs along one dimension *R_COM_*, after aligning the far-ends of two converged FE curves to zero, at which (*i.e.* when *R_COM_* > 50 Å) we assume there is no interaction between any two dimers. Figure 2 presents the free energy profile as a function of the distance between the COMs of two H3 dimers or CENP-A dimers, demonstrating that the FEP minima for (CENP-A/H4)_2_ and (H3/H4)_2_ are appropriately at −7 kcal/mol and −3 kcal/mol. Since the overall FEP curve of CENP-A dimers is deeper, we expect that, in the absence of DNA and other histone proteins, CENP-A/H4 dimers can more readily assemble into a tetramer than H3/H4 dimers. Furthermore, the free energy minimum is located at a distance of ~28 Å between dimers of CENP-A/H4 and at ~32 Å between dimers of H3/H4 (Figure 2), indicating that the thermodynamically favored CENP-A tetramer is more compact than the H3 tetramer. This result agrees quantitatively with previous SAXS measurements that found the CENP-A tetramer to be substantially more compact relative to its H3 counterpart (16). Also, Banks and Gloss used far-UV circular dichroism to measure the folding and unfolding kinetics of (H3/H4)_2_ experimentally (49). The free energy of the dimer-tetramer reaction they obtained is −2.6 kcal/mol. Our computed minimum, at −3 ± 0.2 kcal/mol, is in qualitative agreement with their experimentally measured data. Overall, in agreement with experimental work (Black *et al* (34)), we find that the CENP-A homotypic tetramer, on its own, is thermodynamically more stable, and more compact, than the tetramer of H3/H4. Additional free energy profiles projected on other reaction coordinates, both one-dimensional and two-dimensional, are provided in SI (Figures S4 and S5).

**Figure 2:**
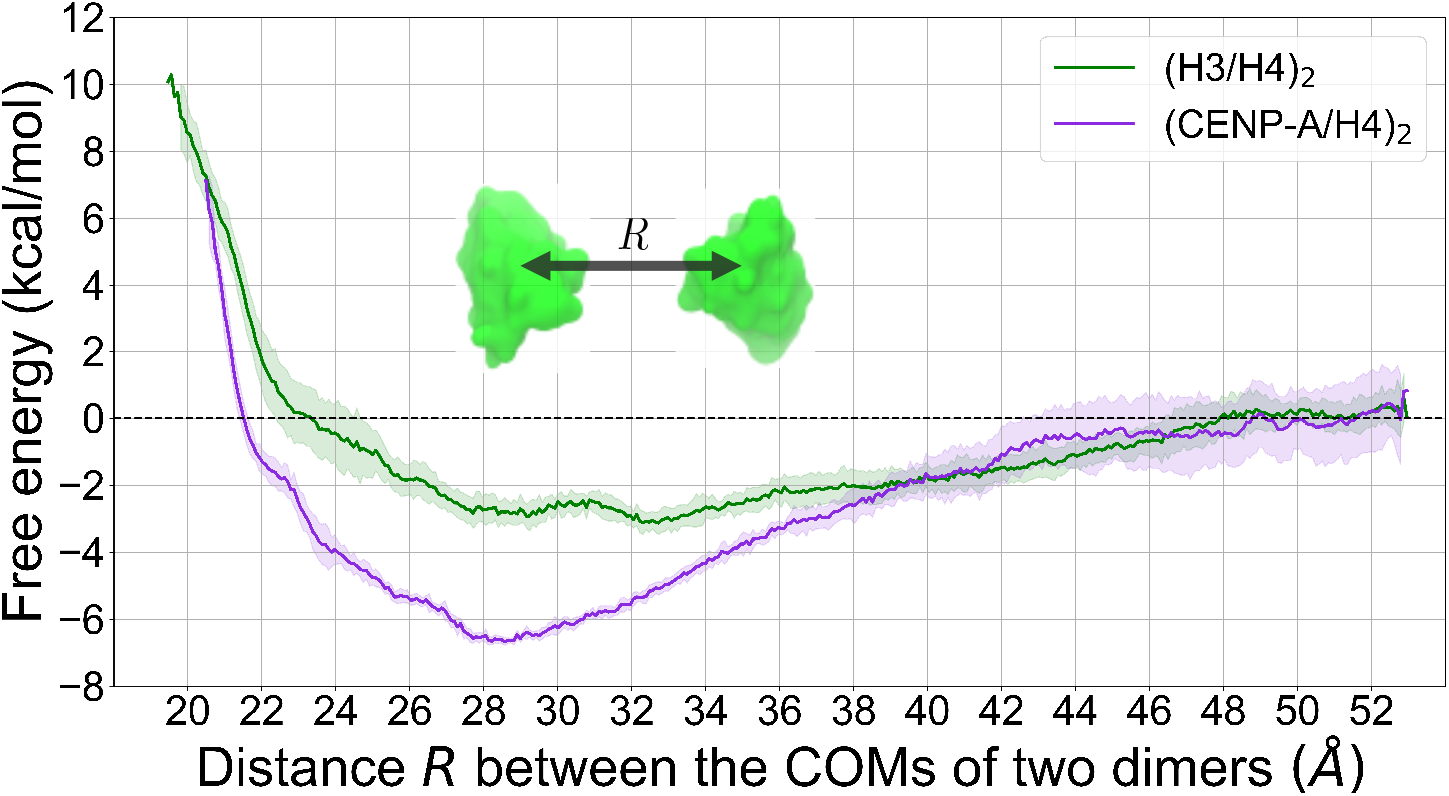
(CENP-A/H4)_2_ has a deeper free energy profile than (H3/H4)_2_. The potential of mean force (PMF) along the distance *R* between histone dimers is deeper for (CENP-A/H4)_2_ (purple) than for (H3/H4)_2_ (green). *R* is measured from the center-of-mass (COM) of one dimer to the other. The shaded areas illustrate the standard deviations of the curves.

### Tetramer geometries and the swiveling dynamics

To further explore the intrinsic dynamics of histone tetramers, we performed microsecond-scale continuous constant temperature CG-AWSEM simulations for CENP-A and H3 tetramers, starting from pre-assembled conformations taken from the central tetramers of the corresponding octameric nucleosome crystal structures (Figure 3A). Other structures from octamer or chaperone-tetramer complexes could have been used as well because the overall structures of the tetramer are nearly identical despite divergent crystallization conditions (Figure S4). Overall, our results obtained from these continuous simulations were broadly consistent with the above enhanced sampling simulations, and they provide important dynamics insights. We present here some of the most salient observations; additional analyses including the principle component analysis (PCA) and other structural quantities including the root-mean-square deviation (RMSD), *R_COM_*, and *Q_interface_* can be found in SI (Figures S9, S7A,B,C).

**Figure 3:**
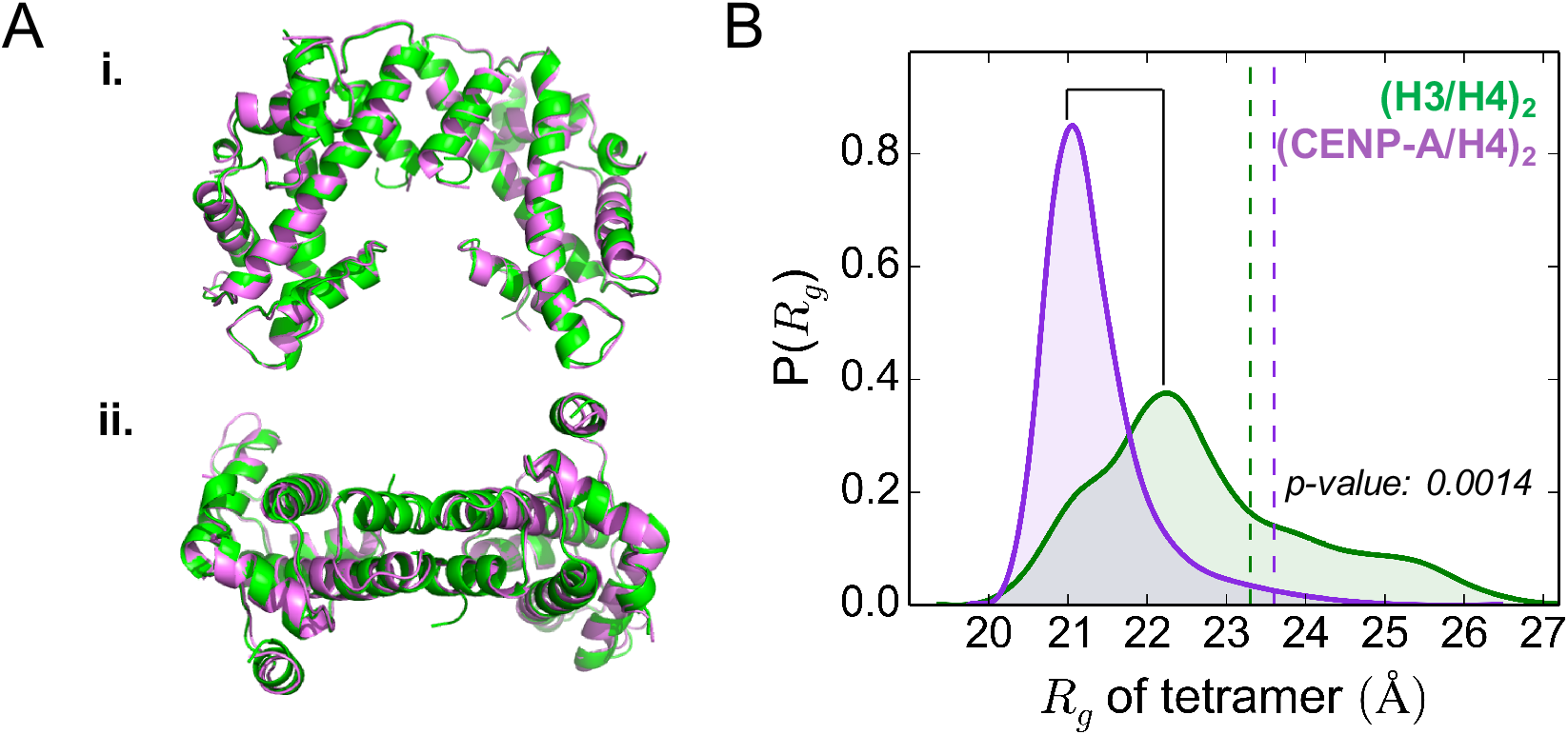
(CENP-A/H4)_2_ is more compact than (H3/H4)_2_. (A) The initial conformations of the H3 tetramer (green) and CENP-A tetramer (purple) were taken from their nucleosome crystal structures (PDB IDs: 1KX5 and 3AN2). Lateral view (i) and top view (ii) of aligned structures are displayed. (B) The CENP-A tetramer has a smaller radius-of-gyration *R_g_* than the H3 tetramer, with a narrower distribution. The vertical dashed lines mark the measured *R_g_* values of the initial structures.

To quantify the molecules’ degree of compaction, we calculated the tetramer’s radius of gyration, defined as 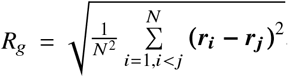, where *N* is the total number of residues and ***r**_i_* are the coordinates of *i*th residue. Figure 3B shows that the average *R_g_* for (CENP-A/H4)_2_ is 21 ± 0.7 Å and 23 ± 1.4 Å for (H3/H4)_2_, implying that (CENP-A/H4)_2_ samples more compact geometries with less *R_g_* fluctuations. The *R_g_* distribution of (CENP-A/H4)_2_ is unimodal, with a dominant central peak, while the H3 tetramer *R_g_* samples a much broader distribution (Figure 3B), consistent with the above free energy calculations (Figure 2). Moreover, the mean value difference between the two systems in our simulation matches the previous experimental data, where Black *et al.* measured that the CENP-A tetramer complex chromatographs as a single species with a Stokes radius of 28 Å, smaller than that of H3/H4, 30.5 Å (34). Together, these results suggest that the CENP-A tetramer is more closely packed, structurally more well-defined than the canonical H3 tetramer.

In recent magnetic tweezer experiments, the DNA of H3-containing tetrasomes were observed to flip between left- and right-handed superhelically-wound states (5, 6), which may be initiated by conformational changes of the proteins inside. To better compare with these experiments, we examined the overall orientation of the simulated tetramers by measuring the dihedral angle between the two composing dimers. We chose to measure the dihedral angle of the two H3 *α*2 helices (similarly for CENP-A), since they are the longest continuous structural component in each dimer molecule.

Our results demonstrate that compared to (CENP-A/H4)_2_, the two H3 dimers in (H3/H4)_2_ occupy a range of orientations, as indicated by the distribution of above-mentioned dihedral angle that includes three populations (Figure 4B): one positive and two negative, three distinct states in total (Figure 4A,B i,ii,iii). Furthermore, (H3/H4)_2_ frequently transits from one dihedral angle to another (fifteen major switches in the measurement of dihedral angle), undergoing swiveling motion around the binding interface (Figure S10 and Supplemental Movie S1). The range of orientations for two histone dimers and its dynamical transition found in our simulations can explain the spontaneous flipping behavior of DNA handedness in the tetrasome as revealed in magnetic tweezer experiments (6). A positive dihedral angle of the tetramer would correspond to left-handed superhelically-wrapped DNA, and vice versa (Figure 4C).

**Figure 4:**
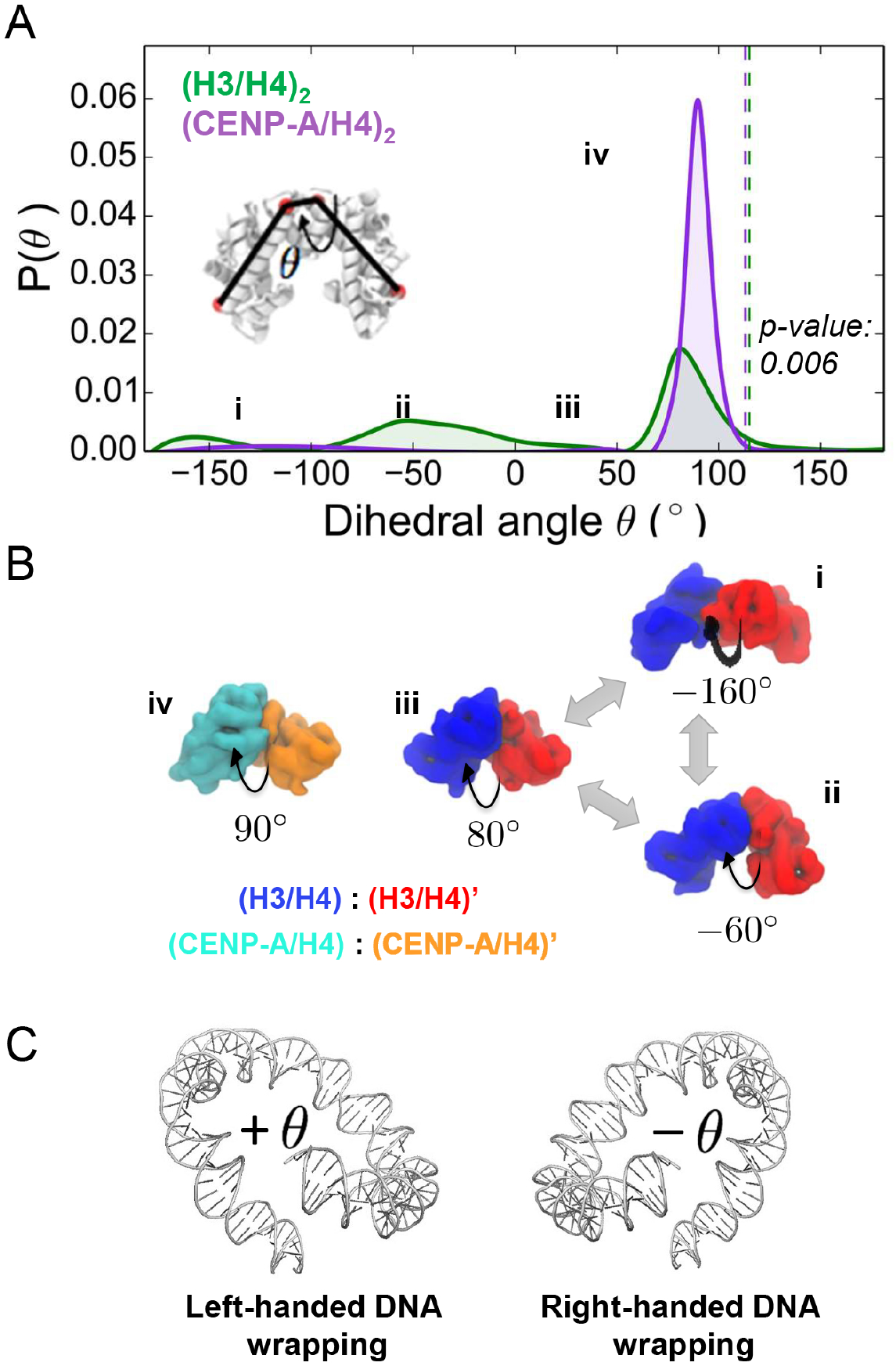
The H3 tetramer swivels around its binding interface while the CENP-A tetramer remains relatively stable. (A) The distribution for the dihedral angle between *α*2 helices features one prominent peak for (CENP-A/H4)_2_, and three smaller peaks for (H3/H4)_2_, indicating (CENP-A/H4)_2_ maintains a more fixed orientation than (H3/H4)_2_. Vertical dashed lines are the corresponding dihedral angles of the initial structures. (B) Representative conformations from each population are displayed. (C) Positive (+) and negative (−) dihedral angles of the histone tetramer measured here correspond to the left-handed and right-handed DNA superhelical wrapping in the tetrasome, respectively.

In contrast, (CENP-A/H4)_2_ maintains a relatively fixed orientation, with no obvious rotational motions around the interface (one switch in the measurement of dihedral angle, Figure S10 and Movie S2). The dihedral angle between the scaffold helices is about 90° (Figure 4A,B iv), less than the angle measured in crystal structures of the CENP-A tetramer from the nucleosome or with chaperones (110°), implying a twisted interface geometry. Indeed, from the simulation snapshots, as well as other measurements including overall *R_g_* and *R_COM_* between dimers, the two CENP-A/H4 dimers seem to pack more closely together in a twisted orientation, presenting a compact tetramer. Moreover, we observe that, in the absence of DNA and other histones, both H3 and CENP-A histone tetramers prefer not to occupy the same plane compared to the geometries of their respective nucleosome structures (Figure 4A). The *α*2 helices of CENP-A were found to be curved (Figure S18) as was also revealed from experimental observation (16). The curvature of *α*2 helices could be a result of the absence of surrounding DNA and bracketing H2A/H2B, which provides necessary topological support to the central tetramer.

### Distinct dynamics at the tetrameric interface

To uncover the mechanistic basis for the observed difference in behavior between CENP-A and H3 tetramers, we then assessed whether it arises from the tetrameric interface (*i.e.* the interface between two dimers). We calculated *Q_interface_* for the continual simulations, referring to the native interface contacts from the crystal structure (PDB: 1KX5). The distribution for the CENP-A tetramer is centered at 0.5, while the same distribution for the H3 tetramer contains three peaks, with an average value of 0.2 (Figure S8B). This result implies that (CENP-A/H4)_2_ forms a tetrameric interface that is better defined and more native-like compared to (H3/H4)_2_. In the context of the DNA-free tetramer, the four-helix bundle of (CENP-A/H4)_2_ which composes the tetrameric interface still maintains a well-connected and symmetric geometric arrangement (Figure 5B), despite some structural twisting. This is not found in the H3 tetramer case.

**Figure 5:**
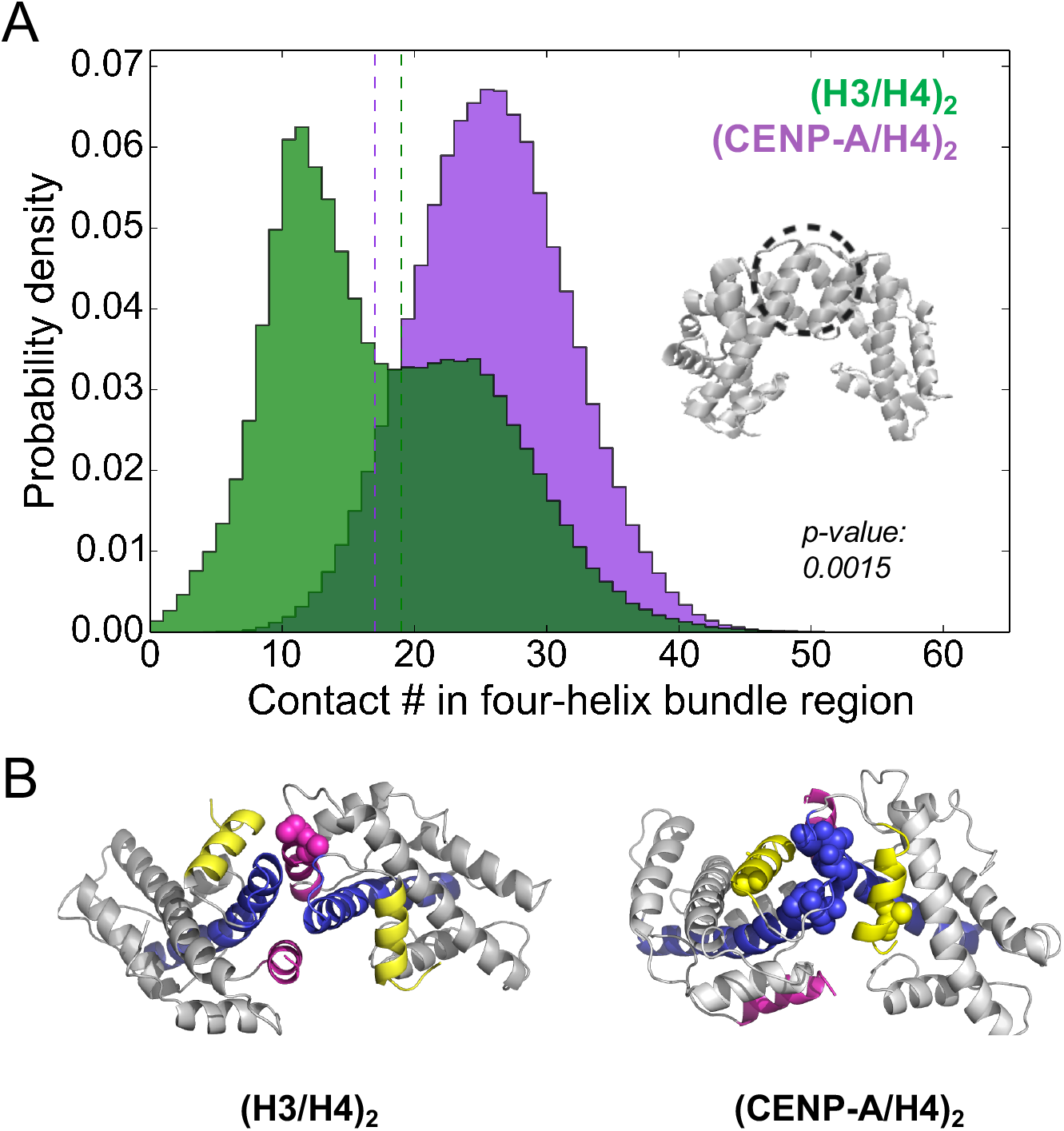
(CENP-A/H4)_2_ has a more stable four-helix bundle than (H3/H4)_2_. (A) (H3/H4)_2_ (green) forms fewer contacts than (CENP-A/H4)_2_ (purple) in the four-helix bundle region. The histogram of the number of contacts for (H3/H4)_2_ has two peaks at 13 and 25 while (CENP-A/H4)_2_ has a single peak at 27. The dash lines mark the four-helix contacts number in corresponding crystal structures. (B) Corresponding representative structures demonstrate that the (H3/H4)_2_ four-helix bundle becomes broken or disrupted by *αN* helices (pink), while the four-helix bundle (*α*2 and *α*3, blue and yellow) remains stable in (CENP-A/H4)_2_ throughout the simulation. *α*2 and *α*3 helices are marked in blue and yellow. CENP-A-specific residues L112, T113, L114, V126 and H3-specific V46 and A47 are shown in coarse-grained spheres.

Furthermore, we performed contact analysis for the four-helix bundle region. It demonstrates that there are more contacts, on average, in the corresponding region of (CENP-A/H4)_2_ (~27) than in the same region of (H3/H4)_2_ (~17) (Figure 5A). Also, one dominant peak is found in the (CENP-A/H4)_2_ contact histogram, but two peaks exist in that of (H3/H4)_2_. Detailed residual pair interactions from AWSEM show that the CENP-A residues Leu111, Gln127, and Arg131 contribute strong hydrophobic interactions to the four-helix bundle tetramer interface (Table S1), which H3 lacks. (CENP-A/H4)_2_ maintains a well-defined, native-like four-helix bundle throughout the simulation (Figure 5B), with the *αN* helices remaining outside the central interface. Note that the previously suggested CENP-A Leu112 residue (16), which is next to Leu111, is not found in the top strong interacting pairs of our simulations. The reason for this discrepancy is unclear.

Meanwhile, we observed structurally heterogeneous H3 *αN* helices, consistent with previous EPR experimental findings (51). Moreover, we notice that the *αN* sections of histone H3 may play an important role in disrupting the four-helix bundle at the H3 tetrameric interface (Figure 5B & S11). Indeed, the distances between the *αN* helices from each H3 copy shows that the H3 *αN* helices feature a considerably wide distribution including two prominent peaks (at about 20 and 32 Å apart) (Figure S9D). Further, this disruption is mainly mediated through the hydrophobic interactions between Val46, Ala47, Leu48 from *αN* and Leu107, Ala111 from *α*2. The αN helix of H3 has greater hydrophobicity than CENP-A does, which could explain, in part, why H3 *αN* helices are more likely to be found close together at the interior of the tetramer than the same helices of CENP-A. We tested this hypothesis by performing similar simulations for both systems but starting from structures excluding *αN* helices. The analyses (Supplemental Section 11) confirmed our hypothesis that the flexible *αN* helices play a significant role in the swiveling motion of H3 tetramer, since switching between different H3 tetrameric dihedrals is significantly diminished when *αN* helices are excluded (Figure S13D). Interestingly, even without *αN* helices, CENP-A still forms more four-helix contacts (Figure S13B) and a more native-like binding interface (Figure S13C) than the H3 tetramer.

Hence, from this analysis we suggest that: (1) specifically in the DNA-free tetramer context, the unique hydrophobic residues (Leu112, Thr113, Leu114, Val126) at the CENP-A:CENP-A interface may help contribute an intrinsically stronger four-helix bundle than H3; (2) the more hydrophobic H3 *αN* helix (Val46, Ala47, Leu48) tends to disrupt the relatively weak four-helix bundle formation and lead to the swiveling motion around the H3 tetramer interface.

### Effects of H2A/H2B on histone tetramers

Finally, we wanted to examine the effects of histone dimer H2A/H2B on the dynamics of tetramers (H3/H4)_2_ and (CENP-A/H4)_2_. We investigated the DNA-free canonical H3 and variant CENP-A octamers using similar simulation procedures. Both the H2A/H2B dimers maintained well-native conformations throughout the simulations (Figure S14D). However, their distances to the central tetramer are diverse for H3 and CENP-A cases (Figure S14C), implying different effects of H2A/H2B on each tetramer.

As done for tetramers, similar analyses such as *R_COM_, R_g_*, and tetrameric dihedral *θ* were performed to explore the dynamics features of the histone octamers. For the H3 octamer, the distribution of both the tetrameric *R_g_* and the distance *R* between H3/H4 pairs becomes more focused and Gaussian-like, compared to the *solo* tetramer situation (*solo* refers to the tetramer in isolation, without any other proteins; Figure 6A vs Figure 3B; Figure S14B vs Figure S9B). The standard deviation decreases from 3.8 Å to 1.9 Å for *R*, and from 1.4 Å to 0.7 Å for *R_g_*, agreeing with previous EPR experimental data (51) showing the reduced H3 tetramer flexibility in an octamer. The distribution of the tetrameric dihedral angles of H3 features a dominant peak at 90° (Figure 6B), similar to that measured in the case of CENP-A, with the other two populations observed in *solo* H3 tetramer simulations diminished. 84% of H3 tetramer conformations in the octamer simulations have a positive tetrameric dihedral angle, significantly more than that in the *solo* tetramer simulations (58%). These data establish that the swiveling motion around the binding interface was reduced due to the bracketing histone dimers H2A/H2B on either side of the tetramer.

**Figure 6:**
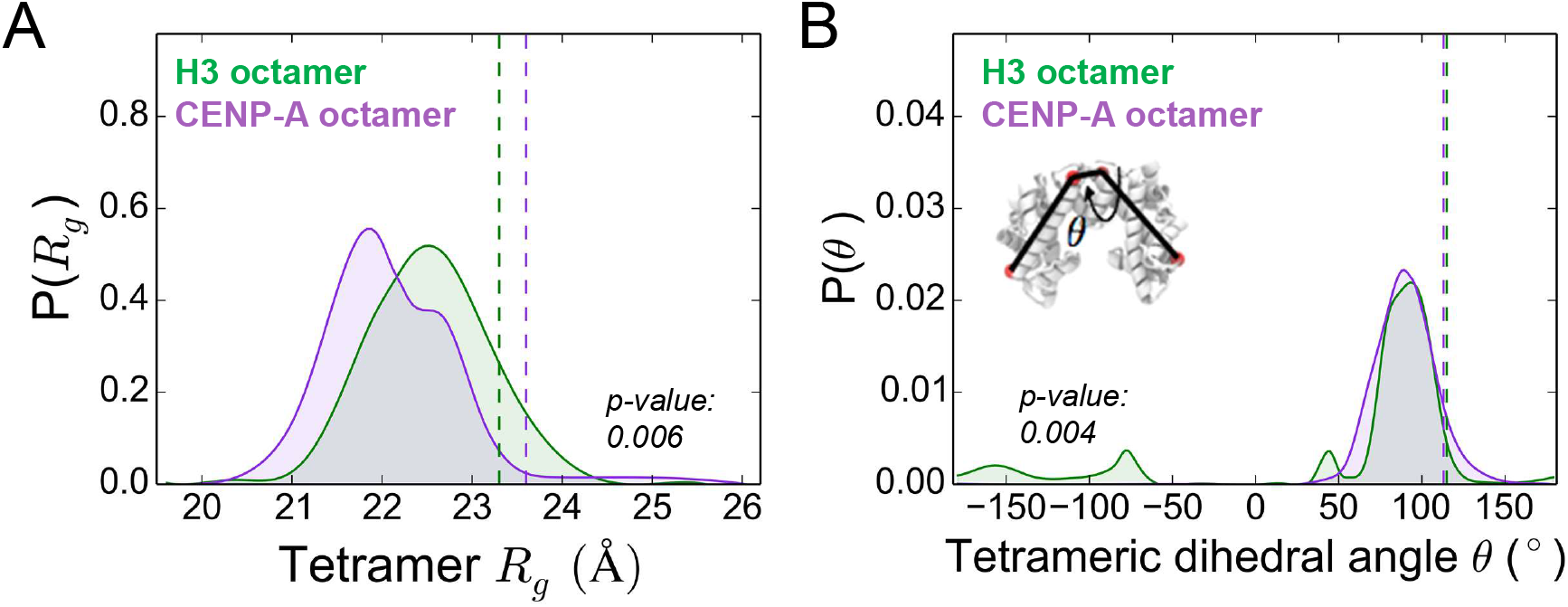
H2A/H2B stabilizes (H3/H4)_2_ but not (CENP-A/H4)_2_. (A) The probability distribution of H3 tetramer *R_g_* features a more focused peak in the context of an octamer compared to that of the *solo* H3 tetramer (Figure 3C), while one peak and one shoulder exist in the same distribution for the CENP-A tetramer in the context of an octamer. (B) Distributions of the dihedral angle between *α*2 helices demonstrate that, in the presence of H2A/H2B, (H3/H4)_2_ becomes more similar to (CENP-A/H4)_2_; both curves feature a prominent peak around 80°. Vertical dashed lines in both panels are the corresponding tetrameric *R_g_* values and dihedral angles measured from the initial octamer.

Nevertheless, analogous stabilizing effects were not found in the CENP-A octamer case. Interestingly, for the CENP-A octamer, a shoulder and a tail are present in the distributions of *R* and *R_g_* of the CENP-A tetramer, indicating new conformational flexibility of (CENP-A/H4)_2_ in the context of an octamer. In particular, the second most populated state has a larger *R_g_* and *R* than the dominant values observed for the *solo* CENP-A tetramer (Figure 6A vs Figure 3B; Figure S14B vs Figure S9B). In turn, this implies that the addition of H2A/H2B dimers leads to a less compact association of CENP-A dimers, encouraging the CENP-A tetramer to adopt a geometry closer to that found in the octameric nucleosome. This frustration between the intrinsic compactness of the *solo* CENP-A tetramer and the expansion and structural twisting induced by the addition of H2A/H2B dimers explains well the observed computational and experimental findings that CENP-A-containing histone nucleosomes or octamers are structurally more flexible and heterogeneous than their canonical counterparts (38, 39).

## DISCUSSION

### Maturation of the nucleosome stepping through dimers, tetramers, and octamers

In this work, we used coarse-grained modeling to study the thermodynamical and dynamical properties of canonical and variant CENP-A histone oligomers. We comprehensively compared the association energy of H3 dimers and CENP-A dimers forming their corresponding homotypic tetramer, inferring an energy difference of 4 kcal/mol between the two types. We also observed that the H3 *αN* helices enhance the lability of H3 tetramer, contributing to the overall swiveling motion around the less hydrophobic H3 tetrameric interface. The addition of (H2A/H2B)s restrains the flexibility of H3 and pushes CENP-A to adopt multiple conformations. Our results are largely in agreement with the prior experimental observations on these systems including H3 tetramer (48, 49) and octamer (51), H3 tetrasome (5) and CENP-A tetramer (16, 34).

We previously reported that in the context of a dimer, histone H4 is more native-like than its binding partner H3 or CENP-A, and that the CENP-A/H4 dimer is more dynamic than its canonical counterpart (37). Here in the context of a tetramer, analyses of the monomer and dimer components yielded consistent results (see Supplemental Section 13). For instance, the average *Q_monomer_* for H4 is larger than that of H3 or CENP-A (Figure S16), implying its noticeable stability; *Q_dimer_* and *Q_dimer,interface_* for H3 are larger, on average, than for CENP-A (Figure S15), indicating that H3 dimers adopt more native-like conformations than CENP-A dimers. However, compared to the structural variabilities within the dimer level, the movements between dimers forming the tetramer are on a larger scale, with an RMSD of 10~15 Å for the tetramer (Figure S9A) versus 3~4 Å for the dimer (Figure S15B, and Figure 2 in ref. 37). Therefore, the dynamics observed here by coarse-grained modeling are unlikely to be sampled ergodically using present-day atomistic simulations.

In our earlier study, the CENP-A nucleosome was shown to be more flexible than the H3 nucleosome, revealing a shearing motion at the tetramer interface (38). Here, in the context of an octamer with H2A/H2B dimers, the CENP-A tetramer occupies two distinct conformational states: one that is similar to that of the isolated tetramer conformation, while the other state is less compact, structurally similar to the H3 (or CENP-A) tetramer in the context of the nucleosomal structure. Therefore, disrupting the energetically stable interface of the CENP-A tetramer likely underpins the shearing motion observed in the octameric CENP-A nucleosome. The two-state memory of the CENP-A tetramer in the octamer may explain why the CENP-A nucleosome is more distortable and dynamic compared with the canonical one.

Decades of work in the chromatin field have demonstrated the crucial importance of not just histone variants, but also the N-terminal tails and post-translational covalent modifications of histones. Together, all of these factors contribute to nucleosome dynamics (59–65) and alter not only the folded state of the chromatin fiber (66–71), but also the affinities of chromatin effector proteins. Additionally, the DNA sequence context is also crucial in determining nucleosome stability (72, 73), nucleosome phasing (74), nucleosome positioning (75–77) and nucleosome spacing (78), all of which determine DNA accessibility (79). These critical epigenetic and genetic components will need to be studied rigorously *in silico* in order to arrive at a holistic representation of the epigenetic landscape of eukaryotic genomes.

### Biological implications

We consider several potential biological implications of our investigation. First, this work emphasizes the importance of structural context for the canonical H3 tetramer, which, *in vivo*, interacts with the surrounding DNA, histone (H2A/H2B)s, or chaperone proteins. The canonical tetramer may have evolved to highly depend on other structural partners, which may be key to ensure the fidelity and stability of genetic material. On the other hand, CENP-A, as a functional variant histone, is intrinsically more stable in its tetramer form, and is therefore less dependent on DNA or other proteins, which may fit better its unique assembly pathway and intricate regulation.

On the basis of our calculations, we speculate that forming the CENP-A tetramer may be nature’s way to reduce the availability of individual CENP-A dimers. The stably formed (CENP-A/H4)_2_ tetramer may serve as a sequestration channel, needed to maintain CENP-A homeostasis. One logical prediction is that histone modification in CENP-A, especially at the interface, which would either strengthen or weaken the rigidity/compactness of the tetramer, might regulate the levels of dimer CENP-A/H4 available for chaperone-mediated assembly to a further extent. On the other hand, we previously found that the disordered terminal tail of CENP-A is very flexible and could easily encounter other proteins (37). The rigidity of the CENP-A tetramer may prevent CENP-A from associating with non-centromeric proteins, so as to avoid the ectopic localization, or promiscuous interactions that might occur more frequently in cancer cells when CENP-A is over-expressed (32).

Another hypothesis based on this research is that the tetramerization of two CENP-A dimers could be nearly irreversible, so that the CENP-A tetramer, once formed, may not be able to separate into two dimers afterwards, even in the presence of chaperone HJURP. In this scenario, the DNA-free protein tetramer might serve as a kinetic trap for excess CENP-A. This hypothesis sheds light on the unique assembly/disassembly pathway of the CENP-A nucleosome. The CENP-A tetramer may be just one state along the assembly pathway of CENP-A nucleosome, after being delivered by HJURP, given the experimental evidence that the CENP-A-CENP-A′ interface is a requirement for stable chromatin incorporation (80).

The CENP-A-specific chaperone HJURP may block CENP-A dimers from self-associating into a tetramer by competing for the same binding site, the internal side of the CENP-A tetramer. It has been shown that two HJURP chaperones (81) and the dimerization of HJURP (29) is required for proper CENP-A nucleosome assembly. Therefore, an HJURP dimer may interact with two isolated CENP-A dimers, instead of with a CENP-A tetramer (Figure 7, right). On the contrary, as in the structure of H3 and its chaperone CAF-1 (82, 83), CAF-1 binds with an H3 dimer at the external side, without the possibility of preventing it from forming a tetramer. Indeed, the kinetically less stable tetramer of H3 may need the enhanced stabilization *via* binding with CAF-1 chaperones at either side (Figure 7, left). Taken all together, we propose two different chaperone models for CENP-A and H3 assembly, CENP-A/H4–(HJURP)_2_–CENP-A/H4 vs CAF-1–(H3/H4)_2_–CAF-1 (Figure 7), with a subtle yet important difference: in the former, two copies of HJURP would prevent two CENP-A dimers from forming a tetramer in pre-assembly complexes, whereas, in the latter, CAF-1 proteins would stabilize a pre-formed H3 tetramer in preparation for subsequent nucleosome assembly. Our results support the possibility that canonical H3- and CENP-A-containing nucleosomes may have orthogonal assembly pathways: while H3 could be deposited as a tetramer, CENP-A may be loaded in the form of a dimer.

**Figure 7:**
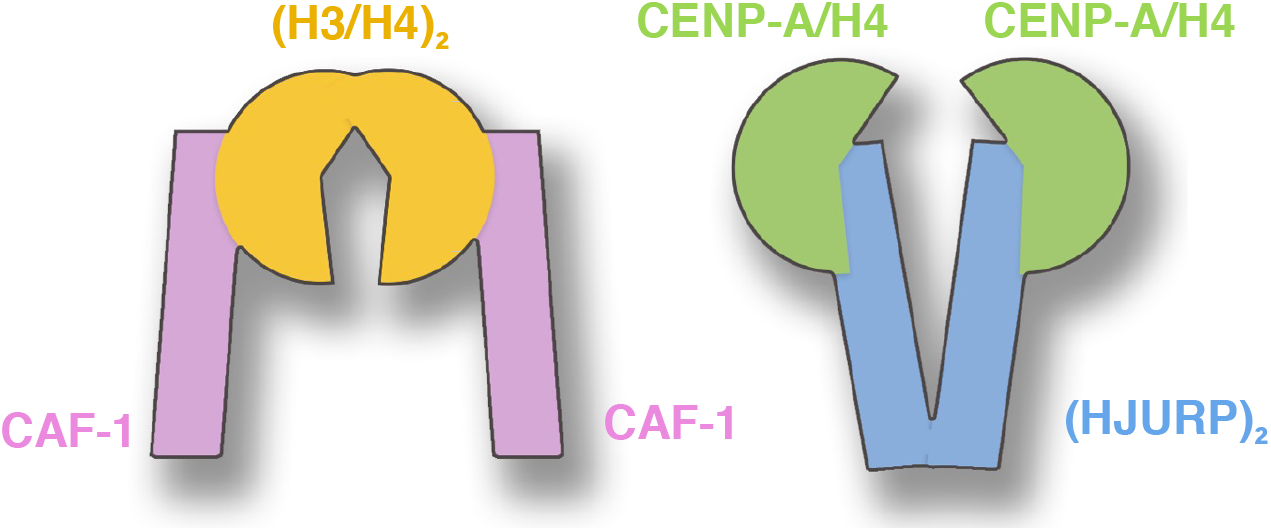
Suggested different models for histones and their chaperones during deposition. (Left) H3/H4 may be deposited in the form of a tetramer with each external side bracketed by a CAF-1 chaperone, which may stabilize the tetramer. (Right) CENP-A may be deposited as dimers; each dimer loaded by one HJURP chaperone.

## CONCLUSION

This work establishes that variant histone CENP-A thermodynamically favors a tetramer formation while the canonical H3 presents remarkable swiveling dynamics about the tetramer interface contributing a rugged yet shallow binding free energy landscape. Moreover, H2A/H2B dimers restrain the internal rotational motion of (H3/H4)_2_ and lead to multiple states for (CENP-A/H4)_2_, providing a possible physical explanation for the shearing motion observed for the CENP-A nucleosome. These findings provide comprehensive molecular insights into the longstanding and recent experimental observations, offering new perspectives on the structural debates over CENP-A dynamics. Based on our results, we suggest two different assembly models for H3 and CENP-A. Lastly, we propose that the (CENP-A/H4)_2_ tetramer may serve as a sequestration channel *in vivo*, which would provide another layer of regulation to ensure the proper homeostasis of CENP-A.

## Supporting information

Movie for H3 Tetramer

Movie for CENP-A Tetramer

## AUTHOR CONTRIBUTIONS

H.Z., Y.D. and G.P. designed the research. H.Z. carried out computational simulations. H.Z. and D.W. analyzed the data. All authors wrote the article.

## ACKNOWLEDGMENTS

H.Z. thanks Drs. Aram Davtyan and Bin Zhang for helpful discussions of AWSEM method, Drs. Minh Bui, Ignacia Echeverria and Longhua Hu for critical reading of the manuscript, and Ms. Sizhu Li for graphics figure. This work was supported by the NSF Grant CHE-1363081, the intramural research program of CCR/NCI, and the NCI-UMD Partnership for Integrative Cancer Research. H.Z. is also supported by the Ann G. Wylie Dissertation Fellowship from University of Maryland, College Park.

## SUPPLEMENTARY MATERIAL

An online supplement to this article is available.

