## Supplemental Information for "The Oligomerization Landscape of Histones"

### Contents


**1 AWSEM Model Details** **3**

**2 Trajectory Analysis Details** **4**

**3 Error Analysis and $P$-value Calculation** **5**

**4 Simulation Convergence** **7**

**5 Umbrella Sampling Histograms** **8**

**6 Crystal Structure Alignments** **9**

**7 Amino Acid Sequences** **11**

**8 Extended 2D/1D Free Energy Profiles from Enhanced Samplings** **12**

**9 Extended Distributions of Structural Measures and Measurement with Time in Constant T Simulations** **15**

**10 Principle Component Analysis in Constant T Simulations** **18**

**11 Simulations of Tetramers Excluding $\alpha N$ Helices** **20**

**12 Extended histone octamer simulations results** **22**

**13 Analyses on the Level of Dimers and Monomers** **23**

**14 AWSEM Energy Analysis of the Tetramer Interface** **26**

**References** **30**

### 1 AWSEM Model Details

The Associative-memory, Water-mediated, Structure and Energy Model (AWSEM) was developed based on decades of efforts. History of related models can be referred to chapter 3 of a recent book by papoian *et al.*[1] It has been successfully applied to study protein folding,[2] binding,[3,4] aggregation,[5–7] intrinsically-disordered proteins,[8] membrane proteins,[9,10] and protein-DNA association.[11–13] Basically it is a transferable coarse-grained protein force field based on the energy landscape theory.

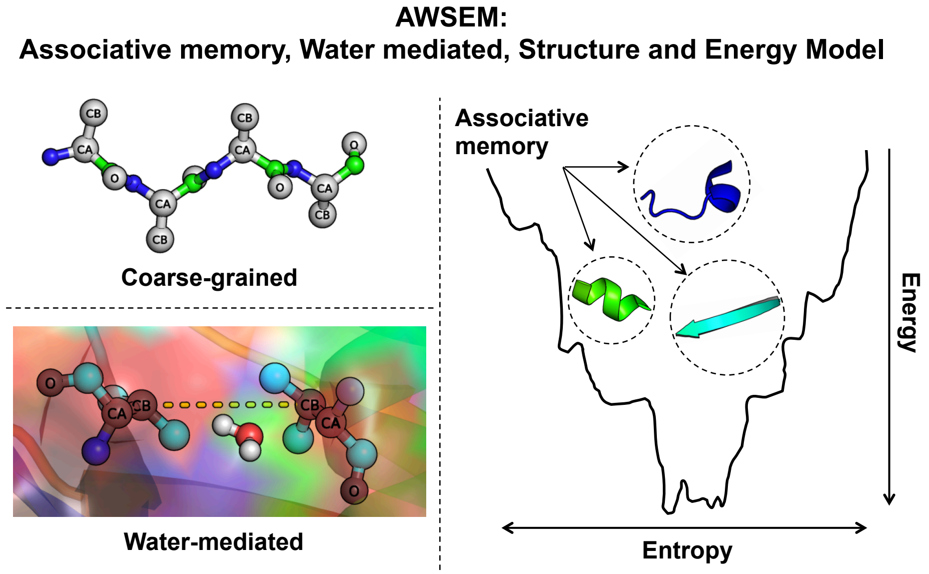


Figure S1: **A cartoon diagram of the Associative memory, Water mediated, Structure and Energy Model (AWSEM)** This diagram shows three main features of this model. Firstly, it uses three beads ($C_\alpha$, $C_\beta$, $O$) to represent one amino acid. Residue information such as charge or radius is carried by $C_\beta$ bead. Secondly, it features a mediated potential for water-residue interactions. Thirdly, it contains a local structure based biasing potential term, aligning the target sequence to short peptides with known structures in the pdb database. Figure adopted from ref. 14

Here, we used the AWSEM model as the force field to perform molecular dynamics simulations. In AWSEM, both physically-motivated potential terms, such as the water-mediated potential, the hydrogen-bonding potential, and a bioinformatically-based local structure biasing term are included. Three beads represent one amino acid and a water-mediated potential describes the water-protein interactions. Written in Eq. S1, the AWSEM Hamiltonian includes a backbone term $V_{backbone}$, a contact term $V_{contact}$, a many-body burial term $V_{burial}$, a hydrogen-bonding term $V_{HB}$, and the bioinformatical term, called the fragment-memory or associative-memory potential $V_{AM}$. The protein-like backbone is maintained by the term $V_{backbone}$, a combination of harmonic potentials based on the positions of $C_\alpha$, $C_\beta$ (H for Glycine) and O atoms. $V_{contact}$ and $V_{burial}$ deal with the water-mediated or protein-mediated residual interactions which are based on the local density of residues and their distances in between each. The $V_{HB}$ term defines hydrogen-bonding networks that are responsible for the formation of $\alpha$ helices or $\beta$ hairpins. The bioinformatic term, called the fragment-memory or associative-memory potential $V_{AM}$ is a Go-like potential, but uses fragments of the target sequence to bias the local structure formation. Detailed function form for each potential term can be found in the SI of ref. 2.

$$V_{AWSEM} = V_{backbone} + V_{contact} + V_{burial} + V_{HB} + V_{AM} \tag{S1}$$

Parameters for this model were optimized self-consistently using energy landscape theory, basically maximizing the ratio of the folding temperature to the glass transition temperature $T_f/T_g$.[15] In the current simulations, single memories, typically 12-residue long, were set, exclusively from the histone monomers found in the corresponding nucleosomal crystal structures. Comparisons and analyses regarding different crystal resources are provided in the following section 6 (Figure S4). The weight of memory term $V_{AM}$ is 0.2. Weights for $V_{contact}$ and $V_{burial}$ are both 1.0. The overall potential weight $\epsilon$ is 1.0.

AWSEM simulations were performed in the large-scale atomic/molecular massively parallel simulator (LAMMPS), using the Nosé-Hoover thermostat with a timestep of 5 fs.

### 2 Trajectory Analysis Details

We use the coordinates of C$\alpha$ to calculate the root-mean-square deviations (RMSD), radius-of-gyration ($R_g$), distances ($R$), dihedral angles $\theta$, $Q$ values, and contact analysis. $Q$ values were calculated separately for the tetramer interface between two dimers, the whole dimer, the dimer interface between two monomers, and also for the monomers. The dihedral angle calculations in Figure 4 were obtained by measuring the dihedral angle formed by the first and last C$\alpha$ atoms of the $\alpha$2 helices. A contact in Figure 5 was considered to exist when the distance between two C$\alpha$ atoms was shorter than 8 Å.

The angle between two $\alpha$ helices was calculated by the orientation vector for each selected helix, based on the coordinates of $C_\alpha$ atoms. We then built the variance matrix $V_{helix}$, composed of all the $C_\alpha$ coordinates and coordinates of the geometric center of the helix. Singular value decomposition (SVD)[16] was applied to determine the eigenvalues of the matrix. The eigenvector corresponding to the biggest eigenvalue provided the orientation vector. The variance matrix $V_{helix}$ was defined as:

$$V_{helix} = \begin{bmatrix} x_1 - x_0 & y_1 - y_0 & z_1 - z_0 \\ x_2 - x_0 & y_2 - y_0 & z_2 - z_0 \\ . & . & . \\ . & . & . \\ x_i - x_0 & y_i - y_0 & z_i - z_0 \\ . & . & . \end{bmatrix},$$

where $(x_i, y_i, z_i)$ represents the position of the $i$th C$\alpha$, and $(x_0, y_0, z_0)$ is the coordinates of the geometric center of the selected helix.

### 3 Error Analysis and $P$-value Calculation

Error analysis for the free energy profiles (FEPs) consisted of two parts (Eq. S2). The first part is more related to the convergence of the simulation data. We determined this part of the statistical errors by calculating the FEP variances from independent simulation interval blocks. For example, in Figure 2, we divided the entire simulation trajectory into 3 non-overlapping blocks along the time series, and calculated the free energy for each block independently. The standard deviation of the free energy for each reaction coordinate window determined from the three blocks were taken to be the statistical error from the ensemble. In Eq. S2, $N$ is 3 and $F_i$ is the FEP of the $i$th internal block. $F_0$ is the FEP of the whole simulation data. The second error part is concerned more with the stochasticity of the data. We estimated this part using the Monte Carlo bootstrap error analysis in WHAM.[17,18] The basic idea is, for each simulation window, to use the computed cumulant of the histogram of the real data to randomly generate a new histogram, with the same number of points and then perform WHAM iterations on the set of newly-generated histograms until it is converged, storing the average normalized probability and free energy for each bin in the histogram. The statistical uncertainty is then obtained accordingly.

$$Error(FEP) = \sqrt{\frac{\sum_{i=1}^{N}(F_i - F_0)^2}{N-1}} + Error_{(WHAM)} \tag{S2}$$

We used the p-value from a t-test[19] to verify whether the differences of our samples were statistically significant. T-statistic for mean is given by $(\mid A_1 - A_2 \mid)/\sqrt{\frac{s_1^2}{n_1} + \frac{s_2^2}{n_2}}$, where $A_1, A_2$ are the mean values of the distributions and $s_1, s_2$ are the standard deviations and $n_1, n_2$ the numbers of the data in each distribution. The same method was used for probability density function figures of the main text and SI.

### 4 Simulation Convergence

We calculated the root-mean-squared inner-product (RMSIP)[20] to verify the convergence of all performed simulations. RMSIP, as defined in the below equation, quantifies the overlap between essential subspaces through the inner product of the first ten principal eigenvectors of C$\alpha$ atom coordinates. It is a normalized parameter, where an RMSIP closer to 1.0 indicates greater overlap between data sets.

$$\mathrm{RMSIP} = \left(\frac{1}{10}\sum_{i=1}^{10}\sum_{j=1}^{10}(\boldsymbol{\eta_i}\cdot\boldsymbol{\nu_j})^2\right)^{1/2} \tag{S3}$$

In our simulations, the RMSIP for every individual run was computed between the data sets corresponding to two halves of increasingly higher percentages of the entire simulation trajectory, starting with the first 10 ns, then the first 20 ns, and so forth. All the RMSIP values are over 0.7, indicating adequate convergence of corresponding simulations.

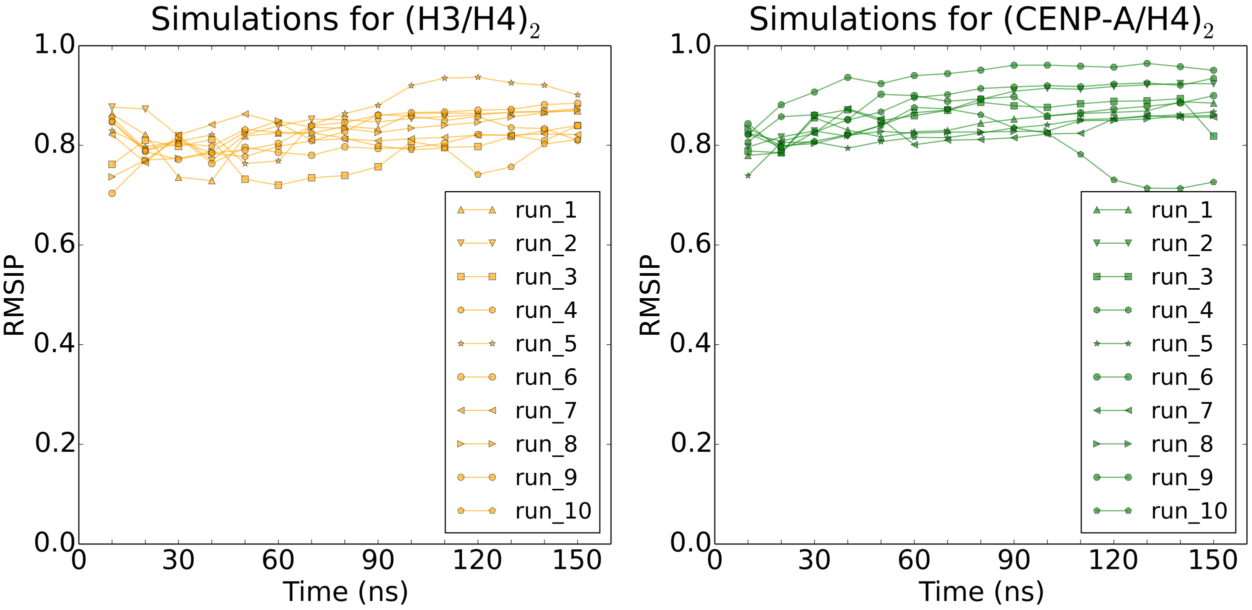


Figure S2: **RMSIP analysis shows the convergence of CG-AWSEM simulations.** RMSIP are calculated for every independent simulation of both H3/H4 and CENP-A/H4 tetramer. All calculated RMSIP are over 0.7, indicating adequate convergences.

### 5 Umbrella Sampling Histograms

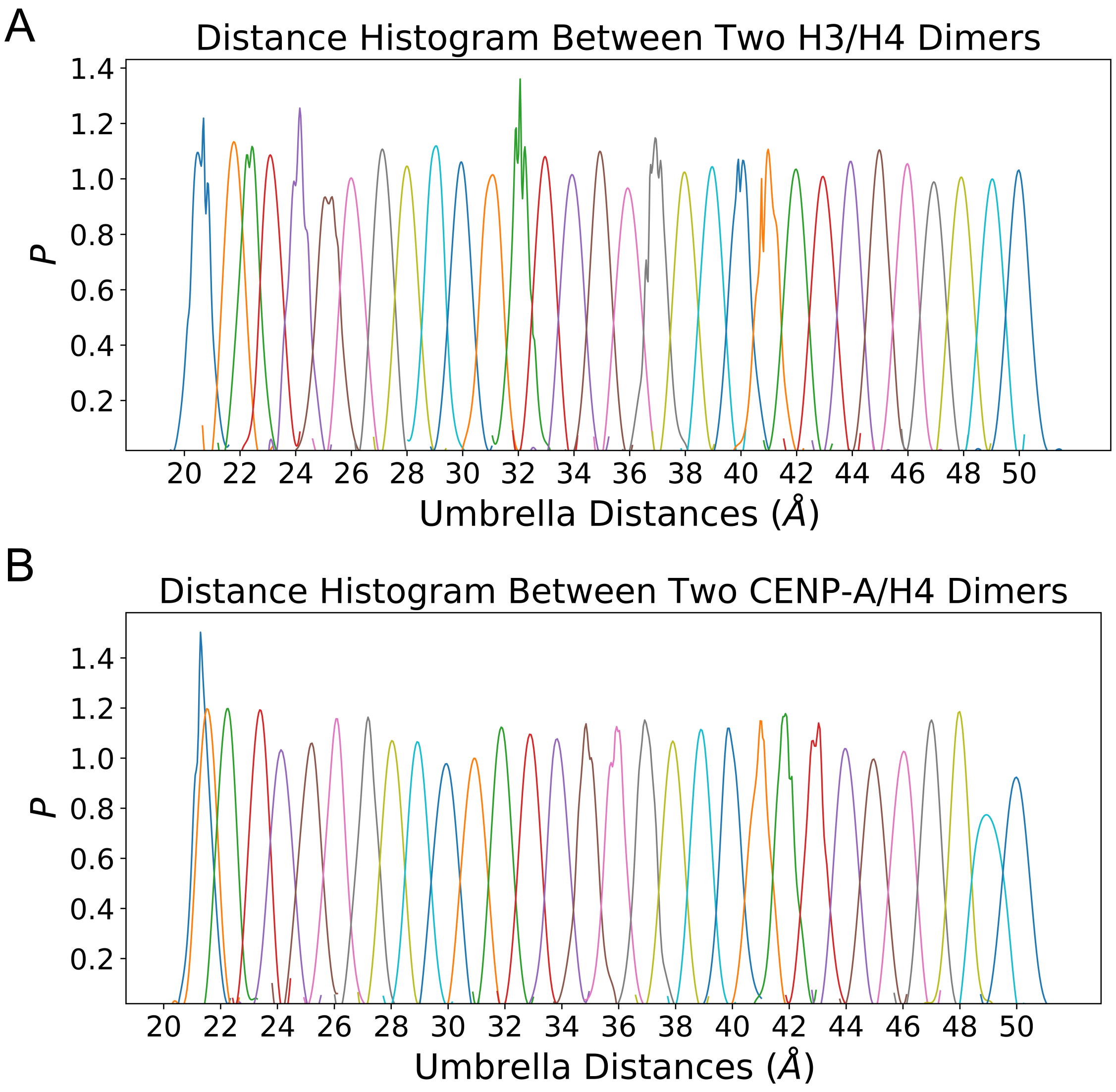


Figure S3: **Sufficient overlaps of reaction coordinate $R$ between adjacent umbrella windows ensure the convergence of WHAM[17] calculation.** Distances from all umbrella windows at replica 300K are collected and histogramed for (A) two H3/H4 dimers and (B) two CENP-A/H4 dimers. PMFs were then calculated from these data using WHAM.

#### 6 Crystal Structure Alignments

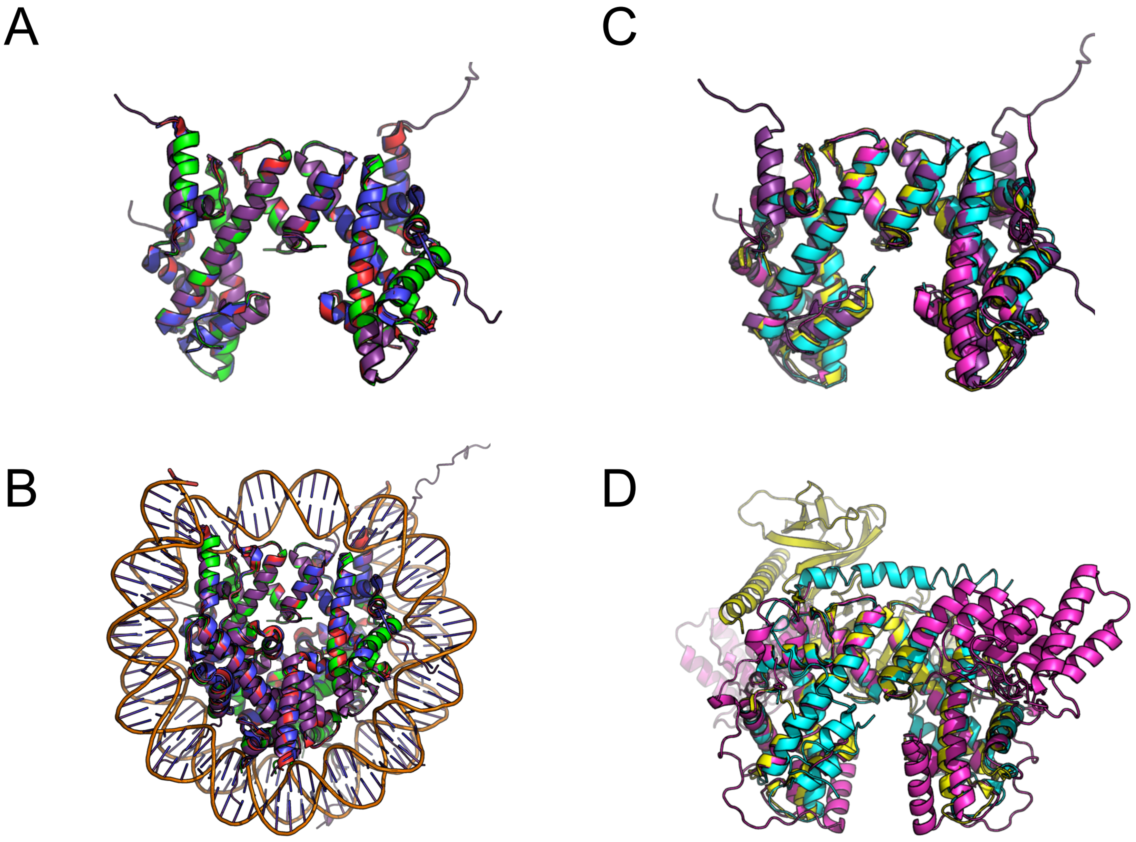


Figure S4: **$(H3/H4)_2$ tetramer presents virtually superposable conformations in the crystal structures of histone octamer, nucleosome and protein complex with chaperone protein(s).** (A) Tetramers from histone octamers and nucleosome are aligned. The average pairwise RMSD is 0.4. Red marks PDB structure 2ARO, blue 2ARO, green 2HIO, and purple is for tetramer from nucleosome structure 1KX5. (B) Structural alignments based on tetramer but shown in the original octamer and nucleosome contexts. Color schemes are the same as in (A). (C) Alignments of histone tetramer from protein complexes that involve tetramer and other proteins. Yellow is for tetramer with chaperone FACT (pdb: 4Z2M), cyan for tetramer with Spt2 (pdb: 5BSA), pink for tetramer with TONSL and MCM2 (pdb: 5JA4), purple refers to histone tetramer from nucleosome (pdb: 1KX5). (D) Same alignments/color schemes as in (C) are shown in their original protein complex environments.

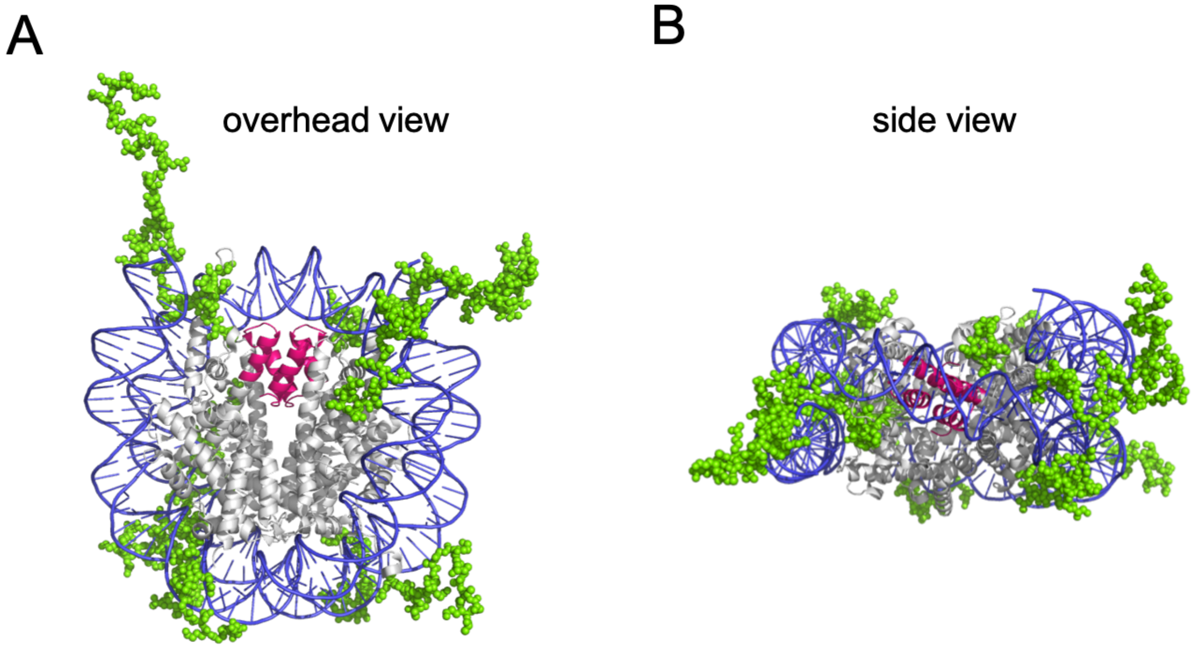


Figure S5: **Crystal structure of nucleosome highlights the position of histone tails and DNA.** Histone tails (green) generally extend out of the histone protein core (grey), closely interacting with DNA (blue) and other potential nucleosome binding partners. The tetramer interface of the histone protein core, four-helix bundle, are marked as red. Shown here is an example of canonical nucleosome structure (PDB: 1KX5), from the overhead view (A) and side view (B).

#### 7 Amino Acid Sequences

**H3 (seq # 46-134):**

VALREIRRYQKSTELLIRKLPFQRLVREIAQDFKTDLRFQSSAVMALQEASEAYLVALFEDTNLCAIHAKRVTIMPKDIQLARRIRGER

**H4 (seq # 25-112):**

NIQGITKPAIRRLARRGGVKRISGLIYEETRGVLKVFLENVIRDAVTYTEHAKRKTVTAMDVVYALKRQGRTLYGFGG

**H3' (seq # 48-135):**

LREIRRYQKSTELLIRKLPFQRLVREIAQDFKTDLRFQSSAVMALQEASEAYLVALFEDTNLCAIHAKRVTIMPKDIQLARRIRGERA

**H4' (seq # 24-112):**

DNIQGITKPAIRRLARRGGVKRISGLIYEETRGVLKVFLENVIRDAVTYTEHAKRKTVTAMDVVYALKRQGRTLYGFGG

**CENP-A (seq # 46-134):**

GWLKEIRKLQKSTHLLIRKLPFSRLAREICVKFTRGVDFNWQAQALLALQEAAEAFLVHLFEDAYLLTLHAGRVTLFPKDVQLARRIRG

**H4 (seq # 25-112):**

NIQGITKPAIRRLARRGGVKRISGLIYEETRGVLKVFLENVIRDAVTYTEHAKRKTVTAMDVVYALKRQGRTLYGFGG

**CENP-A' (seq # 48-135):**

LKEIRKLQKSTHLLIRKLPFSRLAREICVKFTRGVDFNWQAQALLALQEAAEAFLVHLFEDAYLLTLHAGRVTLFPKDVQLARRIRGL

**H4' (seq # 24-112):**

DNIQGITKPAIRRLARRGGVKRISGLIYEETRGVLKVFLENVIRDAVTYTEHAKRKTVTAMDVVYALKRQGRTLYGFGG

**Mark code**: Four-helix Bundle H3 or CENP-A $\alpha 2$ helix

Figure S6: **The sequence number and sequence alignments for the histone proteins investigated in this study.** The amino acid sequences of H3, H4, and CENP-A, provide the primary level of description for the protein structures of the (CENP-A/H4)$_2$ and (H3/H4)$_2$ tetramers. Sequences of the four-helix bundle region are marked in red and sequences of the $\alpha 2$ helix are underlined.

### 8 Extended 2D/1D Free Energy Profiles from Enhanced Samplings

Through calculating the unbiased probability distribution and re-histogramming over different collective variables, we projected the calculated free energy profile onto different coordinates, either two-dimensional or one-dimensional. All the results consistently demonstrate that the H3 histone tetramer occupies a more rugged free energy landscape while CENP-A has a well-funneled landscape topology, indicating that the CENP-A tetramer favors a stable thermodynamic state while H3 does not. The one-dimensional free energy profiles from the coupled replica-exchange and umbrella sampling method can be qualitatively compared to the probability distribution of the same coordinate (Figure 3, Figure S9) from the later long-timescale constant temperature simulations, after using the Boltzmann relation. The consistency between the two results proves the efficiency and convergence of both methods.

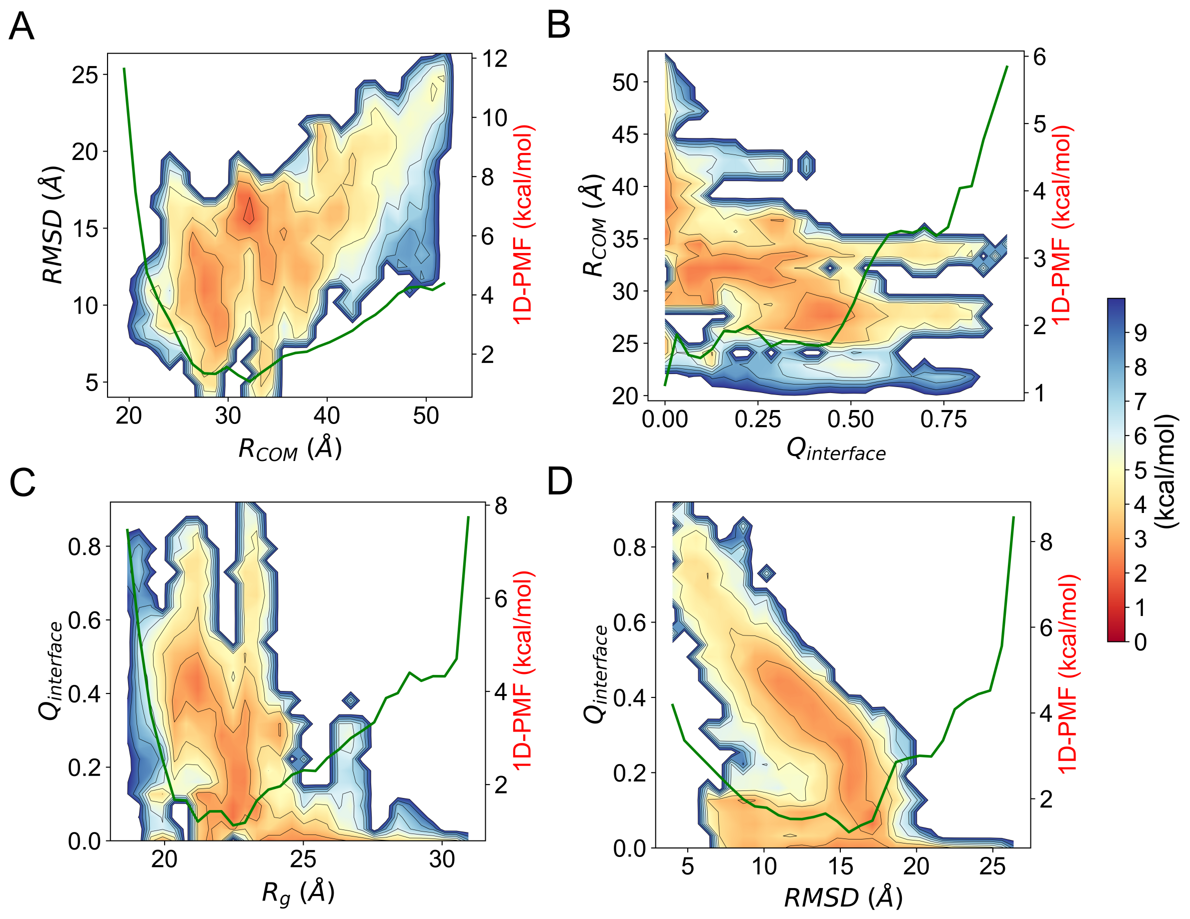


Figure S7: **Extended 2D and 1D free energy profiles for histone $(H3/H4)_2$.** Free energy profiles calculated from the enhanced sampling for $(H3/H4)_2$ tetramer are projected on the 2D reaction coordinates of the distance between centers of masses of each dimer $R_{COM}$ and the measure of overall structural fluctuation root-mean-square-deviation RMSD (A), the tetramer interface contact quantification parameter $Q_{interface}$ value and $R_{COM}$ (B), the geometry measurement radius of gyration $R_g$ and $Q_{interface}$ (C), the RMSD and $Q_{interface}$ (D). 1D free energy projection on the dimension (marked as green line) of $R_{COM}$, $Q_{interface}$, $R_g$, and RMSD are shown on the right side of each panel of (A, B, C, D) accordingly.

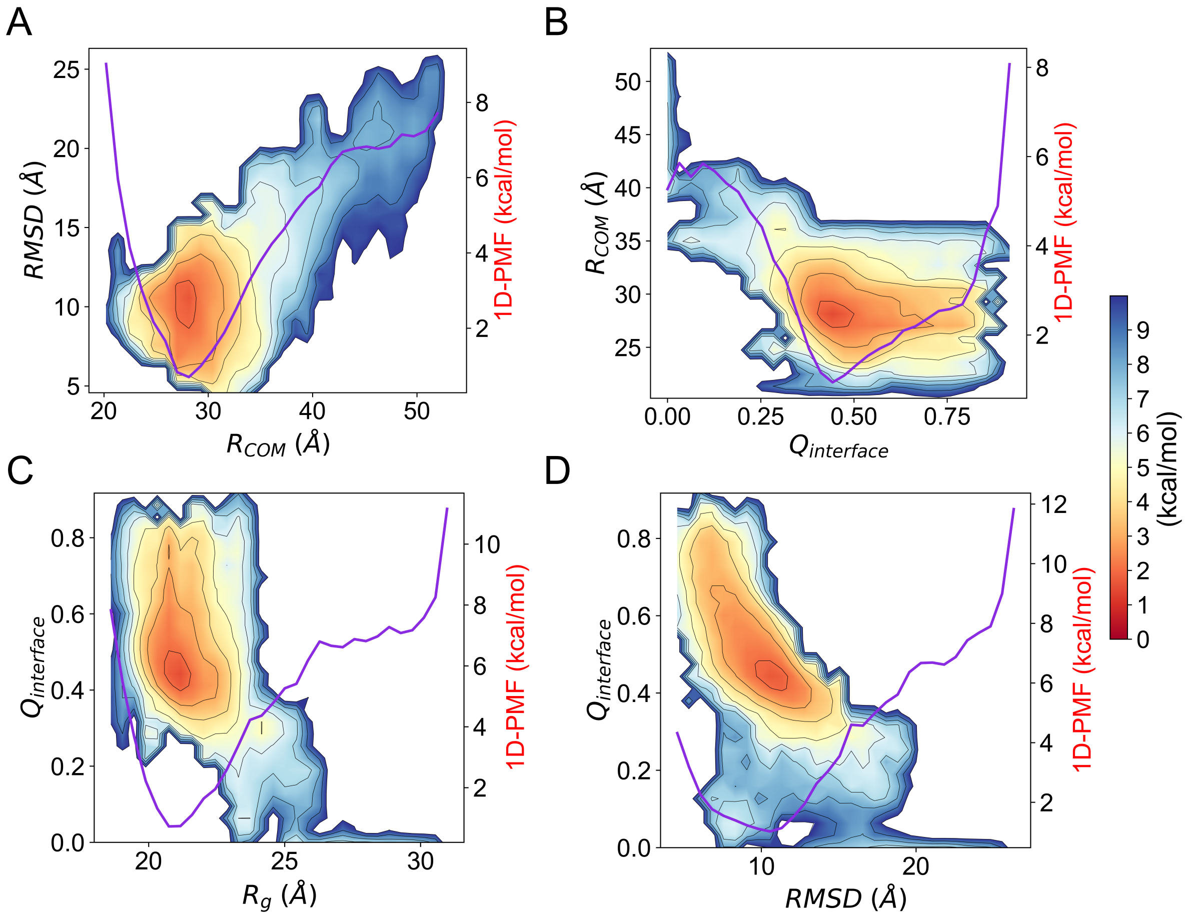


Figure S8: **Extended 2D and 1D free energy profiles for (CENP-A/H4)$_2$ histone tetramer.** Free energy profiles calculated from the enhanced sampling for CENP-A tetramer are projected on the 2D reaction coordinates of the distance between centers of masses of each dimer $R_{COM}$ and the measure of overall structural fluctuation root-mean-square-deviation RMSD (A), the tetramer interface contact quantification parameter $Q_{interface}$ value and $R_{COM}$ (B), the geometry measurement radius-of-gyration $R_g$ and $Q_{interface}$ (C), the RMSD and $Q_{interface}$ (D). 1D free energy projection on the dimension (marked as purple line) of $R_{COM}$, $Q_{interface}$, $R_g$, and RMSD are shown on the right side of each panel of (A, B, C, D) accordingly.

### 9 Extended Distributions of Structural Measures and Measurement with Time in Constant T Simulations

For the long-timescale constant temperature simulations, we also calculated the probability distribution for different structural measures, including the root-mean-square deviation (RMSD), the distance between two internal dimers $R_{COM}$, and the interface contact resemblance $Q_{interface}$. Locations of the distribution peaks observed from constant T simulations agree with the minima locations in free energy profiles from the enhanced sampling (Figures 1&2, Figures S9), demonstrating the convergence of both methods.

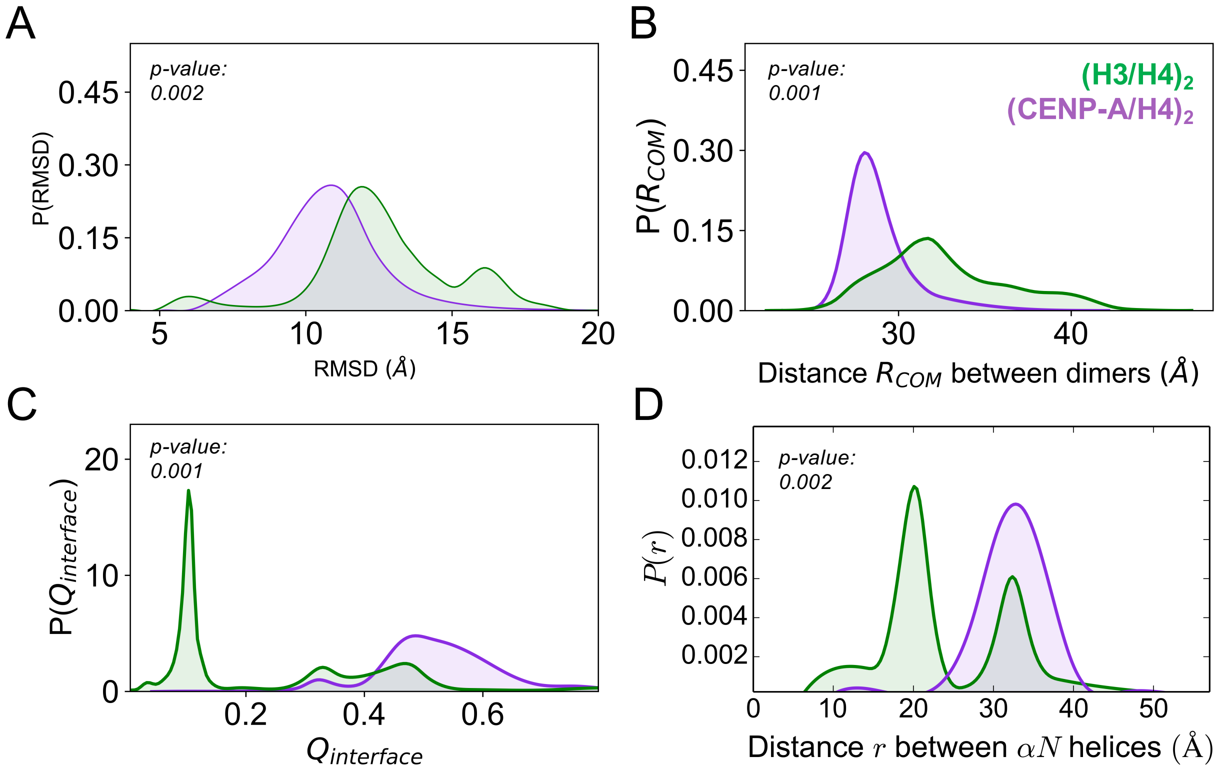


Figure S9: **Distributions of different structural measures confirm the conformational heterogeneity of the H3 tetramer (green) and the homogeneity of the CENP-A tetramer (purple).** (A) The RMSD distribution features multiple populations for the H3 tetramer and a single population for the CENP-A tetramer. (B) The distance between dimers $R_{COM}$ is shorter on average for (CENP-A/H4)$_2$, with a much narrower distribution, than that of (H3/H4)$_2$. (C) $Q_{interface}$ distributions indicate that the interface of (CENP-A/H4)$_2$ remains more stable and closer to its native state than (H3/H4)$_2$. Locations of the peaks in these panels agree with the minima locations in free energy profiles calculated from the enhanced sampling simulations. (D) $\alpha N$ helices of H3 exhibits large heterogeneity while CENP-A $\alpha N$ helices stay stable. The distance measurement between H3 $\alpha N$ helices has two peaks (green) indicating their instability and relocation while the same measurement for CENP-A has one single peak (purple) implying a stable behavior.

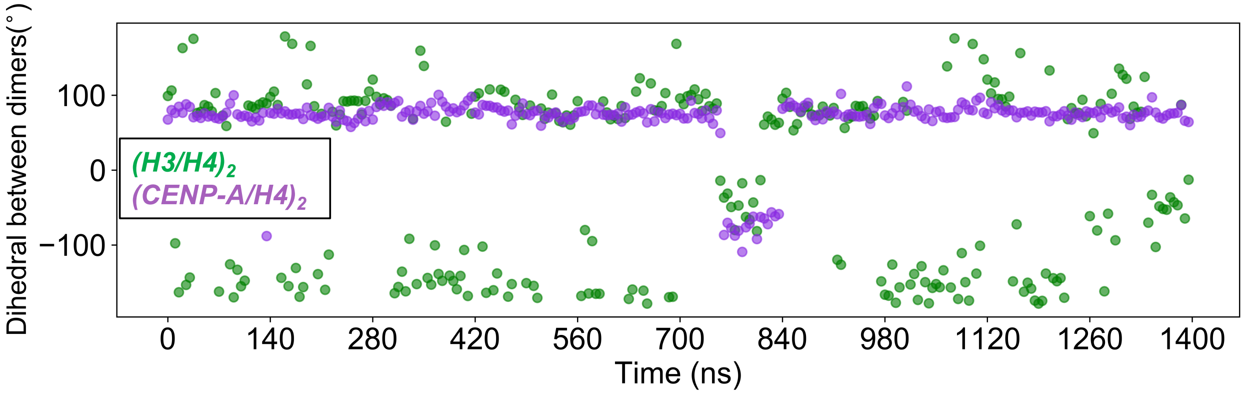


Figure S10: **The dihedral angle between $\alpha 2$ helices measured as a function of time emphasizes the rotational dynamics of the H3 tetramer.** The tetramer dihedral angle of H3 (green) frequently transitions between 90°, −150°, and −50°, while the dihedral of CENP-A (purple) remains constant throughout most of simulation, with only one dihedral angle transition observed.

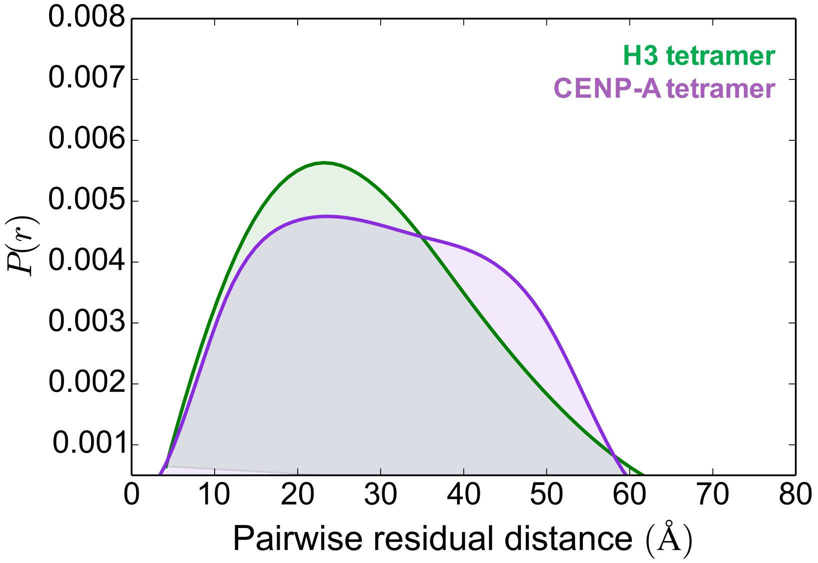


Figure S11: **The distribution of pairwise distance of $C_\alpha$ in representative structure of H3 and CENP-A tetramer.**

#### 10 Principle Component Analysis in Constant T Simulations

To extract the dominant modes of motion from the long-timescale constant temperature MD simulations, we performed principal component analysis (PCA). Overall translational and rotational motion of the MD trajectories were eliminated by a translation to the average geometric center and by alignment to the energy-minimized structure. Then the simulation trajectories were projected onto the first two principal components to illustrate the corresponding free energies. The result (Figure S12) is qualitatively consistent with the free energy profile computed from the enhanced the sampling (Figure 1, Figures S7&S8).

The PCA method is described in detail below. Using the Cartesian coordinates of all $n$ C$\alpha$ atoms over $N$ simulation snapshots ($t_i$ represents an individual time), we created a trajectory position matrix $\mathbf{Q}$,

$$q_i = (x_1, y_1, z_1, ..., x_n, y_n, z_n), \mathbf{Q} = (q(t_1), q(t_2), ..., q(t_N)) \tag{S4}$$

$$Q_{ij} = q_i(t_j) \tag{S5}$$

From this trajectory matrix $\mathbf{Q}$, we constructed a covariance matrix $\mathbf{C}$. Let $N$ be the number of snapshots, $n$ the number of C$\alpha$s, and $\mathbf{Q}$ ($3n \times N$) be the trajectory position matrix. Hence, we have the covariance matrix $\mathbf{C}$ ($3n \times 3n$) defined in Eq. S6. We then diagonalize the covariance matrix $\mathbf{C}$,

$$C_{j,k} = \frac{1}{N-1} \sum_{i=1}^{N} (Q_{ji} - \langle Q_j \rangle)(Q_{ki} - \langle Q_k \rangle). \tag{S6}$$

$$\mathbf{CM} = \mathbf{M}\Lambda \tag{S7}$$

$m_i$, the columns of $\mathbf{M}$, are orthonormal eigenvectors representing the principal components, and the diagonal values along $\Lambda_{ii}$ are the eigenvalues associated with each principal component. We arranged the eigenvalues from highest to lowest, meaning the first principal component captures the most variance within our dataset, the second principal component captures the second most variance, and so forth. Next, we projected the trajectory matrix $\mathbf{Q}$ onto the first 2 principal components, the eigenvectors corresponding to the 2 highest eigenvalues:

$$\nu_1 = \mathbf{Q} \cdot m_1, \qquad \nu_2 = \mathbf{Q} \cdot m_2. \tag{S8}$$

We then separated the $(\nu_1, \nu_2)$ space covered equally into a grid and obtain joint probabilities, $P(\nu_1, \nu_2)$ for each box within the grid. Finally, the free energy landscape was projected along the first two principal components:

$$F(\nu_1, \nu_2) = -k_B T \ln P(\nu_1, \nu_2) - F_{\min}. \tag{S9}$$

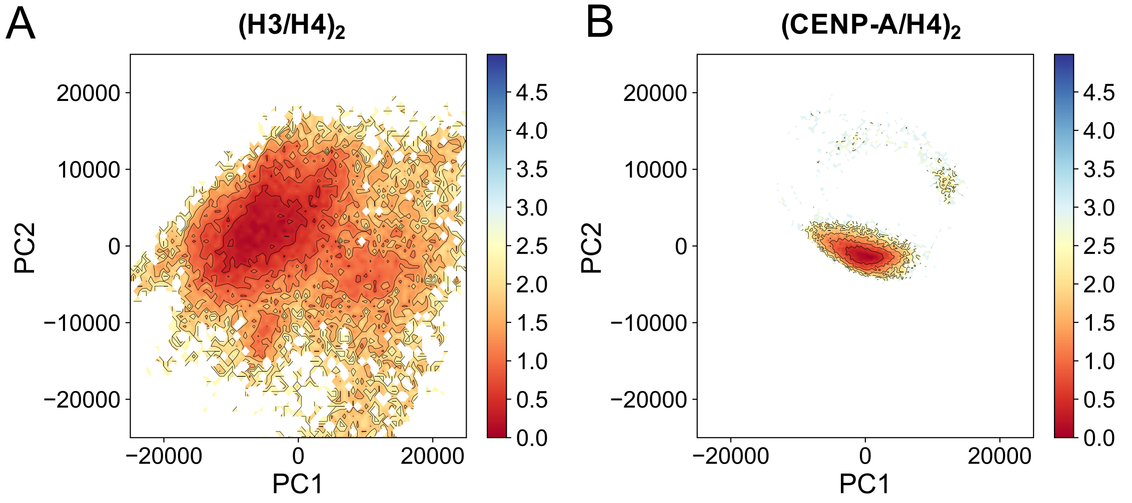


Figure S12: **Free energy projections along the first two PCA components show that $(H3/H4)_2$ occupies a more rugged free energy landscape than (CENP-A/H4)$_2$.** (A) Free energy projection of $(H3/H4)_2$ reveals a broad landscape with multiple conformations basins. (B) Free energy projection of (CENP-A/H4)$_2$ has only one single and deep basin.

### 11 Simulations of Tetramers Excluding $\alpha N$ Helices

Here we performed similar simulations for H3 and CENP-A tetramers excluding $\alpha N$ helices (Figure S13A). It shows that the CENP-A tetramer has a more closely interacting four-helix bundle (Figure S13B), a more native-like interface (Figure S13C), and a more compact formation (Figure S13D), than the H3 tetramer, even when $\alpha N$ helices are excluded, consistent with the SAXS experimental results measured in solution using the same length of proteins.[21] This set of simulations also confirmed our hypothesis that $\alpha N$ is playing a dominant role in leading the swiveling motion around the interface. It can be seen the swiveling between different tetrameric dihedral angles became much less when $\alpha N$ helices were removed (Figure S13D), compared to when $\alpha N$ helices were included (Figure 4A).

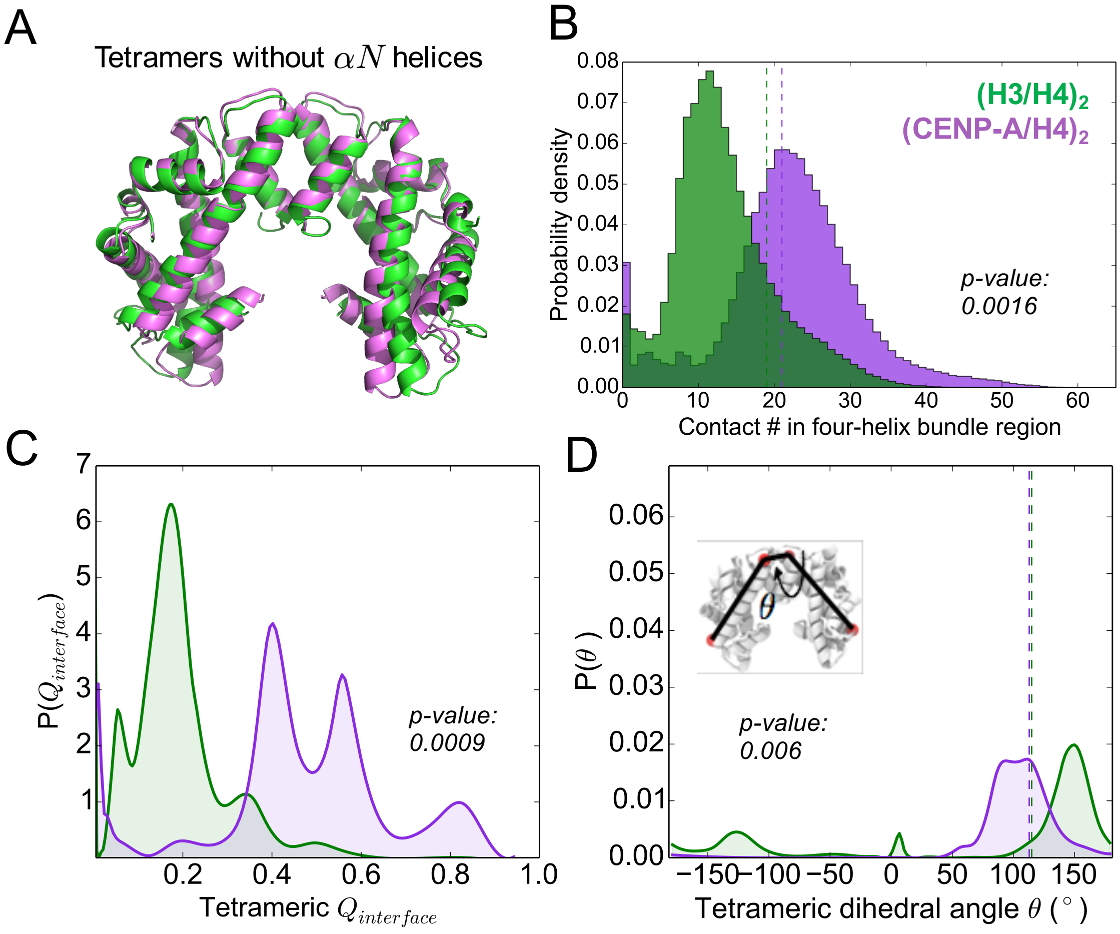


Figure S13: **Tetramer Simulations Excluding $\alpha N$ Helices.** (A) Initial conformations are from $\alpha N$-helix-truncated CENP-A tetramer crystal structure (PDB: 3NQJ, purple) and the same length of H3 tetramer from the nucleosome (PDB: 1KX5, green). (B) More four-helix contacts are found in CENP-A tetramer. The dash lines mark the contacts number in PDB structures. (C) CENP-A has a larger tetrameric $Q_{interface}$, on average than that of H3, indicating a more native-like tetramer interface. The vertical dashed lines are the tetrameric dihedral angle measurements from the nucleosome crystal structures. (D) Tetrameric dihedral angle measurements for CENP-A and H3 tetramer display that CENP-A has a smaller dihedral angle than H3, implying a more compact conformation.

### 12 Extended histone octamer simulations results

Histone octamer simulations start from the octamer formations in the H3 and CENP-A nucleosomes (Figure S14A). Extended octamer simulations analyses show that, (H2A/H2B)s maintained well native-like conformations (Figure S14D) but the interactions between (H2A/H2B)s and the CENP-A tetramer appear to fluctuate more than H3 (Figure S14C).

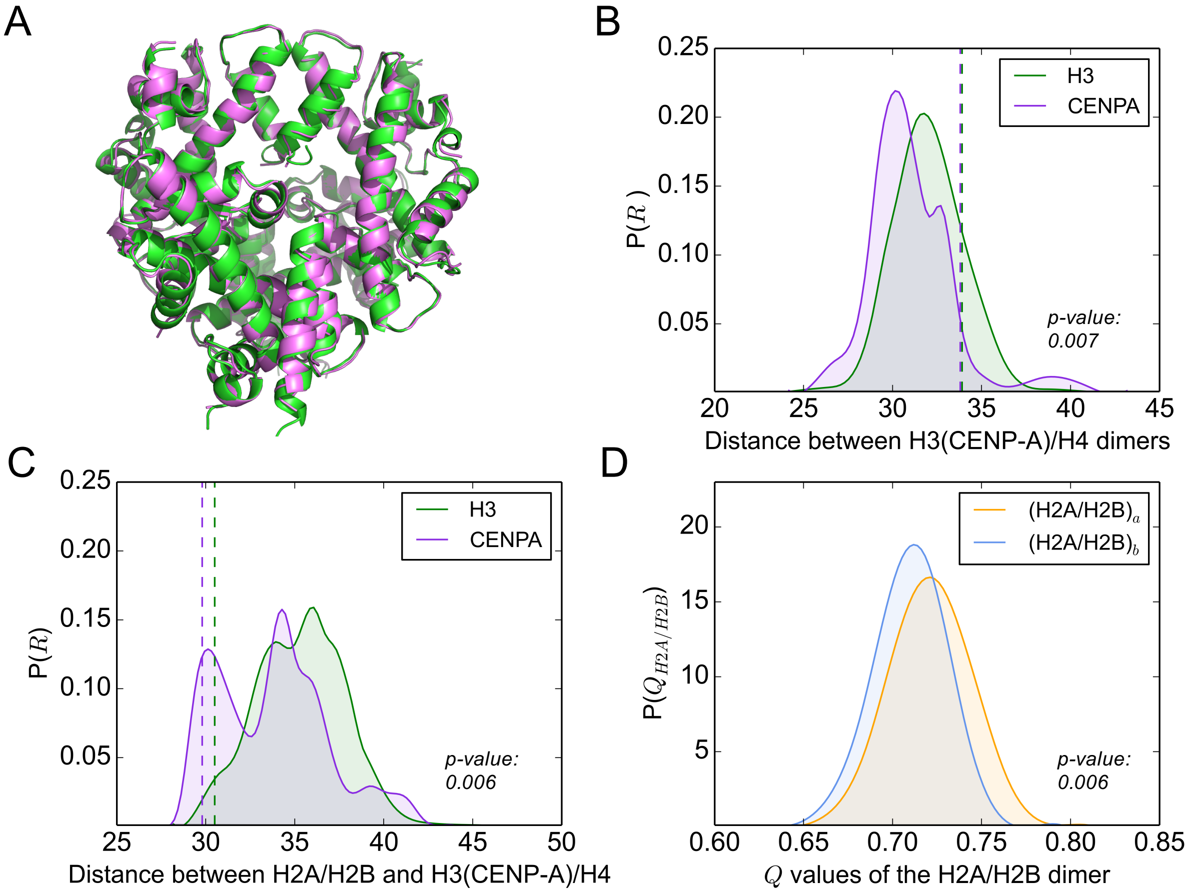


Figure S14: **Simulation results for H3 and CENP-A octamer.** (A) Initial configurations are respectively from the H3- and CENP-A-containing nucleosome structures (PDB: 1KX5 and 3AN2). (B) $R$ distribution shows a single-peak distribution with less deviation for H3 tetramer while CENP-A features one peak and one shoulder in the same distribution. (C)The average distance between (H2A/H2B)s and CENP-A tetramer (purple) is more widely distributed than that of H3 (green). Vertical dashed lines in (B,C) panels are the corresponding distance measurements from the nucleosome crystal structures. (D) $Q$ calculations of (H2A/H2B)s confirm that throughout the octamer simulations, (H2A/H2B)s maintained native-like formations as in their nucleosome structures as both the dimers has high $Q$ values.

### 13 Analyses on the Level of Dimers and Monomers

To compare the results with our previously-published independent dimer study,[4] we performed similar analysis of the tetramer simulations, including RMSD, $Q$ of dimer, $Q_{interface}$ of dimer, and $Q$ for histone monomers. The results here show that, even in tetramer, the CENP-A dimer is still more heterogeneous than the H3 dimer, and the H4 monomer is more native-like than its binding partner H3 or CENP-A.

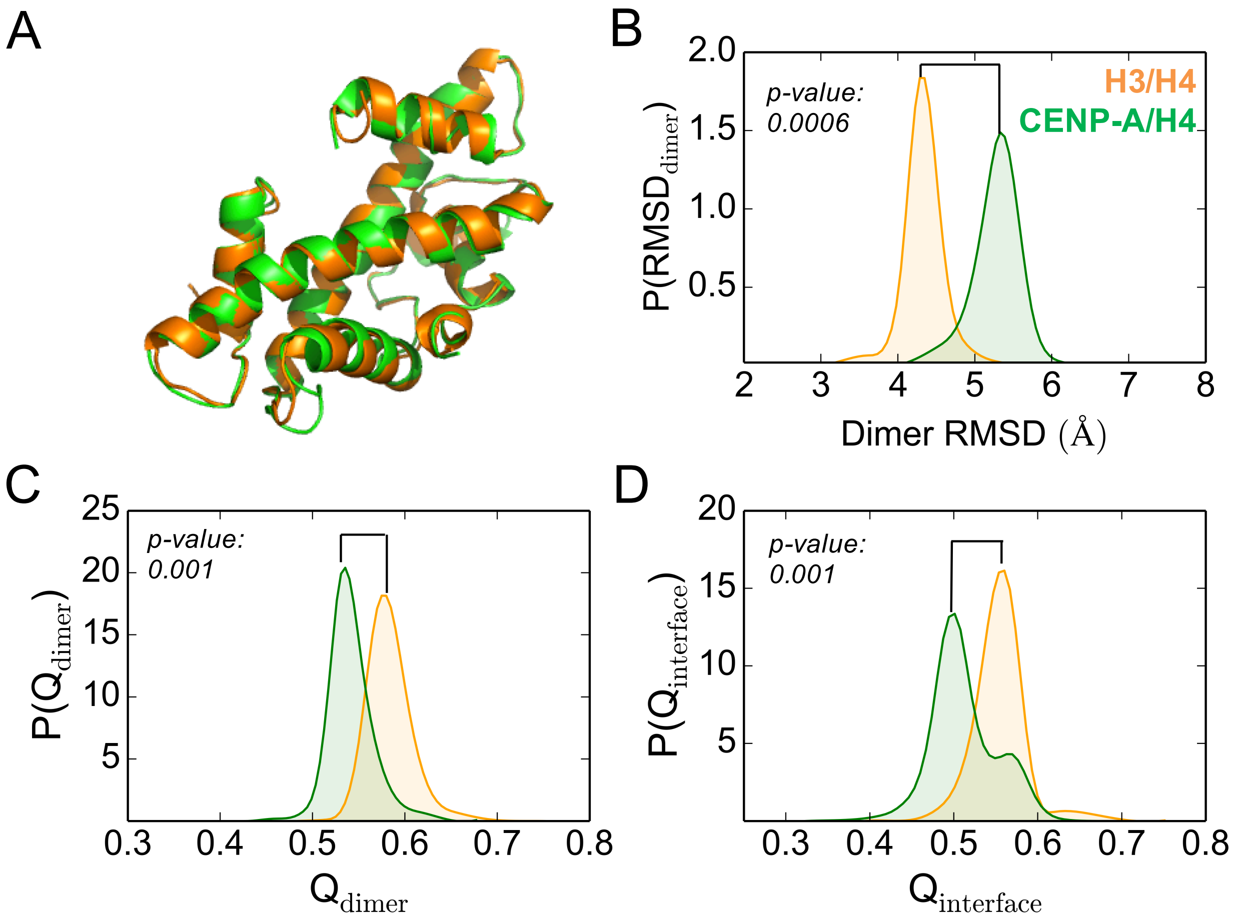


Figure S15: **The H3/H4 dimer is structurally more stable than the CENP-A/H4 dimer in tetramer simulations.** $Q_{dimer}$ and $Q_{interface}$ of the dimer, characterize a dimer's overall structural resemblance or the resemblance of the monomeric interface to its native state (A) respectively. Analyses on the dimer level demonstrate that, in tetramer simulations, CENP-A/H4 exhibits a larger root-mean-square deviation (RMSD) and lower $Q_{dimer}$ (C) and $Q_{interface}$ (D), on average, than H3/H4. This implies the high variability or elasticity of CENP-A in general, which agrees with previous experimental[22,23] and computational studies.[4,24]

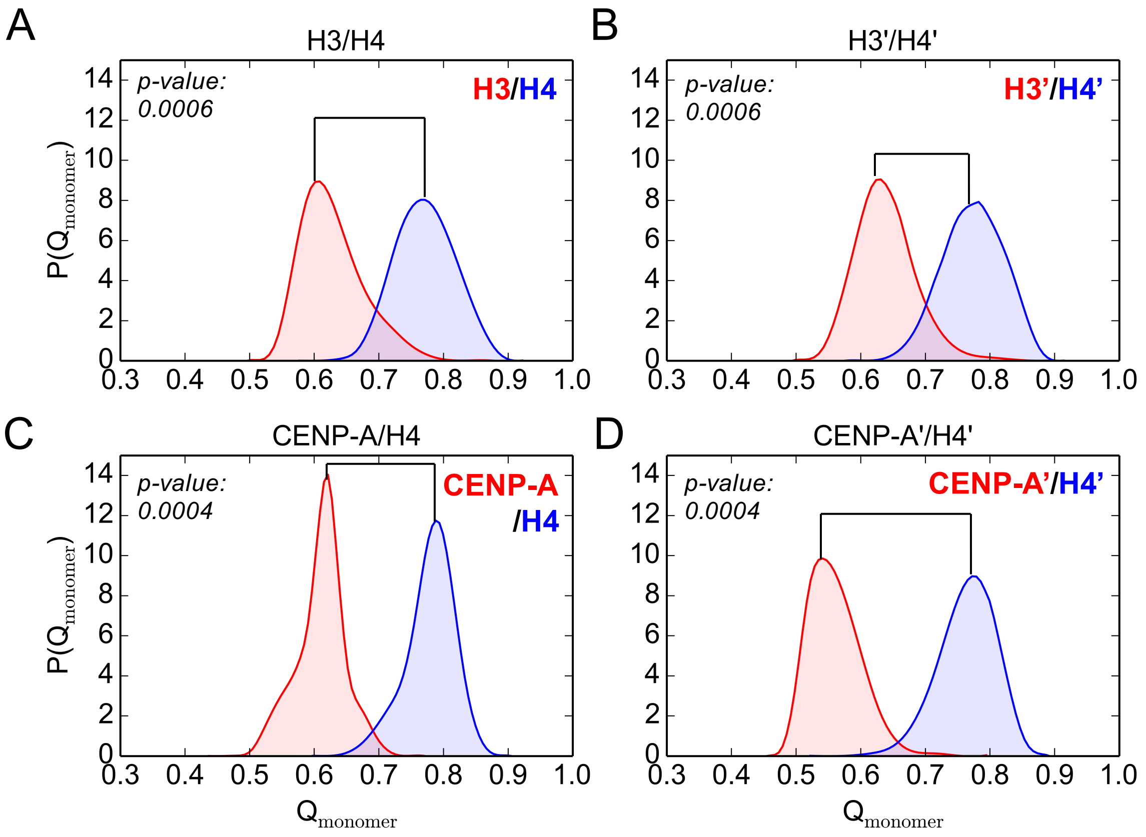


Figure S16: **The H4 monomer maintains a more native-like conformation than its binding partner, either H3 or CENP-A, in all tetramer AWSEM simulations.** $Q_{monomer}$ describes a histone monomer's overall structural similarity with respect to the crystal structures of the corresponding H3 nucleosome (PDB ID: 1KX5) and CENP-A nucleosome (PDB ID: 3AN2). $Q_{monomer}$ was calculated for individual histone proteins of both the first and second H3/H4 dimer (A, B), and for both the first and second CENP-A/H4 dimer (C, D). It shows that H4 has a higher $Q_{monomer}$ than H3, or CENP-A, meaning that H4 maintains a more stable and native-like structure. This result is in accordance with the previous histone dimer study.[4]

#### 14 AWSEM Energy Analysis of the Tetramer Interface

For a better understanding of the histone tetramer interface dynamics at the residual level, we analyzed the energy terms in AWSEM for the corresponding interfaces. The sum of $V_{contact}$ and $V_{bural}$ terms (details in section S1), $E_{pair}$, is collected for every pair of residue interactions between the two dimers inside a tetramer. The sorted large pair interactions are shown in tables S1 and S2. The cutoff was chosen as 0.65 kcal/mol for the absolute value of $E_{pair}$.

The shown two tables (tables S1, S2) are the AWSEM energy outputs for residual pair interactions around the tetramer interface. The first table provides the energies for representative conformations from AWSEM simulations. The energies included in the second table are the AWSEM outputs for the snapshot of each system that is closest to the initial conformation. Both of these two tables demonstrate that CENP-A:CENP-A' has more interacting residue pairs at the binding interface. Detailed residue positions are shown in the structure figures (Figure S17 for H3 and S18 for CENP-A).

Table S1: **Key residue interactions of the tetramer interface in AWSEM (representative conformations)**

| H3 | H3' | $E_{pair}$ (kcal/mol) |
|---|---|---|
| A111 | A111 | -0.9720 |
| N108 | A111 | -0.8886 |
| R116 | A111 | -0.6897 |
| A111 | R116 | -0.6885 |

| CENP-A | CENP-A' | $E_{pair}$ (kcal/mol) |
|---|---|---|
| Y110 | L111 | -1.0000 |
| L111 | T113 | -1.0000 |
| L111 | R131 | -1.0000 |
| T113 | R131 | -1.0000 |
| Q127 | L111 | -1.0000 |
| Y110 | Q127 | -0.9998 |
| T113 | Q127 | -0.9997 |
| T113 | L111 | -0.9784 |
| Q127 | Y110 | -0.9773 |
| R131 | Y110 | -0.9759 |
| Y110 | R131 | -0.9755 |
| R131 | T113 | -0.9072 |
| L111 | Q127 | -0.9071 |
| L111 | L111 | -0.906 |
| L111 | Y110 | -0.897 |
| R131 | A109 | -0.7634 |
| Y110 | Y110 | -0.7006 |
| H115 | H115 | -0.6895 |

Table S2: **Key residual interactions of the tetramer interface in AWSEM (initial conformations)**

| H3 | H3' | $E_{pair}$ (kcal/mol) |
|---|---|---|
| Q125 | R129 | -1.0000 |
| R129 | Q125 | -1.0000 |
| L126 | R129 | -0.9999 |
| R129 | L126 | -0.9999 |
| Q125 | Q125 | -0.9918 |
| N108 | R129 | -0.9832 |
| Q125 | A111 | -0.9549 |
| A111 | Q125 | -0.9263 |
| N108 | Q125 | -0.8416 |
| H113 | H133 | -0.7130 |

| CENP-A | CENP-A' | $E_{pair}$ (kcal/mol) |
|---|---|---|
| Y110 | R131 | -1.0000 |
| L111 | R131 | -1.0000 |
| Q127 | L111 | -1.0000 |
| Q127 | R131 | -1.0000 |
| R131 | L111 | -0.9999 |
| R131 | Q127 | -0.9994 |
| L111 | L111 | -0.9991 |
| L111 | Q127 | -0.9989 |
| L128 | R131 | -0.9878 |
| Q127 | Y110 | -0.9747 |
| R131 | R131 | -0.9349 |
| Y110 | Q127 | -0.8929 |
| L111 | Y110 | -0.8346 |
| R131 | Y110 | -0.6405 |

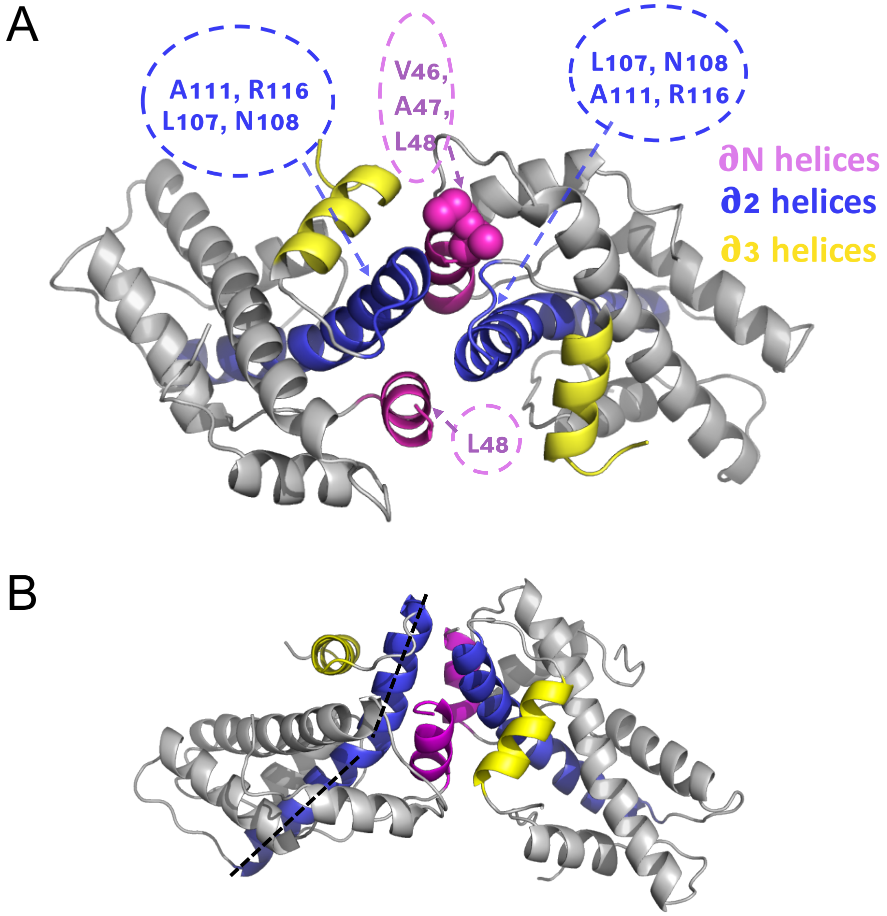


Figure S17: **Representative structure of H3 tetramer with interface interaction details.** (A) Top view of the structure highlights $(H3/H4)_2$ interface is a disrupted *four-helix* bundle region. $\alpha N$ helix competes with the $\alpha 3$ helix, to interact with $\alpha 2$ helix, forming hydrophobic interactions between $\alpha N$ V46, A47, L48 and $\alpha 2$ A111, L107, among which V46 and A47 are H3-specific residues, shown in coarse-grained spheres.(B) Side view of the representative H3 tetramer shows that the $\alpha 2$ helix in H3 can be curved, which is illustrated by the dash line.

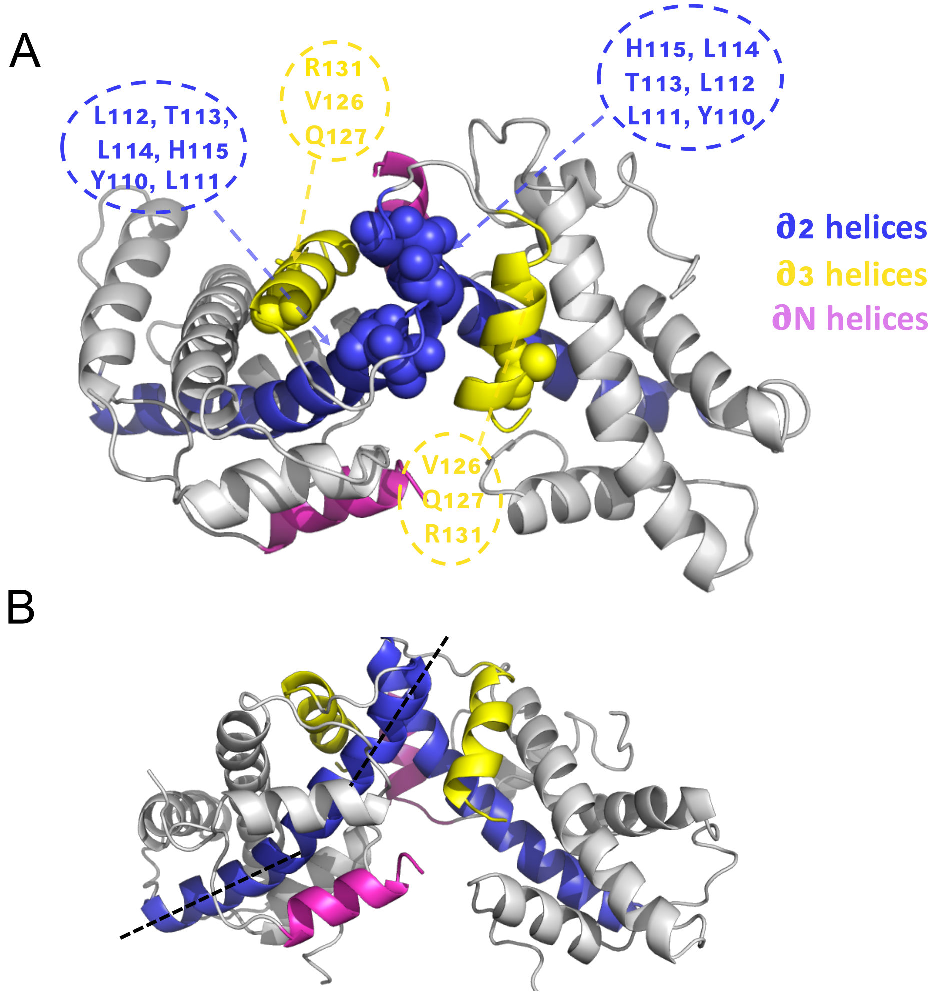


Figure S18: **Representative structure of CENP-A tetramer with interface interaction details.** (A) Top view of the interface highlights a well-formed *four-helix* bundle region, consisting of CENP-A's $\alpha2$ and $\alpha3$ helices. CENP-A-specific residues L112, T113, L114, V126 are shown in coarse-grained spheres. (B) Side view of the representative CENP-A tetramer shows that the $\alpha2$ helix in CENP-A is curved, illustrated by the dash line.
